# Membrane-resolved epithelial electrophysiology revealed using extracellular electrochemical impedance spectroscopy (EEIS)

**DOI:** 10.64898/2026.05.08.723773

**Authors:** Athena J. Chien, Erica Lull, Guiying Cui, Hanna Khor, Nael A. McCarty, Craig R. Forest

## Abstract

Conventional extracellular epithelial electrophysiology measurements report only bulk transepithelial resistance and capacitance, obscuring the distinct electrical properties of the apical and basolateral membranes. This limitation hinders research of epithelial diseases where dysfunction originates at a specific membrane domain—apical or basolateral—for example in cystic fibrosis or toxin-mediated airway injury. Here we present the extracellular electrochemical impedance spectroscopy (EEIS) technique that extracts membrane-specific electrophysiology by fitting impedance spectra to a two-resistor, two-capacitor (RCRC) model. Using human bronchiolar epithelial monolayers (16HBE), we show a correlation between the electrical time constants of the circuit (***τ*_1_ = *R*_1_** · ***C*_1_**, ***τ*_2_ = *R*_2_** · ***C*_2_**) and changes in ion permeability of the basolateral and apical membranes. Experimentally, we show that blocking with 5–10 µM GlyH-101 (i.e. decreasing apical membrane permeability), after 10 µM forskolin activation elicits dose dependent ***τ*_2_** responses that are over 50% larger than ***τ*_1_**and 6–7 minutes faster, whereas 10 µM nystatin (i.e. increasing basolateral membrane permeability) produces ***τ*_1_** responses 21–25% larger than ***τ*_2_** and approximately 2 minutes faster. For cystic fibrosis epithelia, we find that elexacaftor/tezacaftor/ivacaftor (ETI) restores the apical membrane electrical response, resulting in a significant 84% higher ***τ*_2_** than ***τ*_1_** within the first 10 minutes. It also exhibits a greater than 8 min faster ***τ*_2_** response relative to ***τ*_1_** following 10 µM GlyH-101 blocking (i.e., decreasing apical membrane permeability). These results demonstrate that EEIS enables rapid, quantitative, and biologically relevant measurement of apical and basolateral membrane properties in 16HBE epithelia. By providing membrane-specific resolution without the experimental challenges of intracellular electrodes, EEIS establishes a general framework for rapid, membrane-resolved electrophysiology with implications for therapeutic screening.

## 1 Introduction

Epithelial tissues serve as important barriers between various body compartments, often separating organ lumens and the internal environment to maintain organ homeostasis and function [1, 2]. Epithelial cell layers act as a selective barrier to ions, nutrients, and waste products, with transport partitioned across the apical, basolateral, and paracellular pathways—each governed by distinct channel, transporter, and junctional complexes that tightly regulate directional flux and maintain tissue homeostasis (Fig. 1a,i) [1, 3]. Dysfunction of apical and basolateral transport routes contributes directly to disease pathogenesis; for example, in cystic fibrosis, defective apical chloride channel, CFTR, alters transepithelial ion gradients, disrupts mucociliary clearance, and drives chronic airway inflammation [4]. In pertussis, also known as whooping cough, the adenylate cyclase toxin from *Bordetella pertussis* translocates almost exclusively across the basolateral membrane of airway epithelia, triggering cAMP-driven cytotoxicity and barrier dysfunction [5].

**Figure 1:**
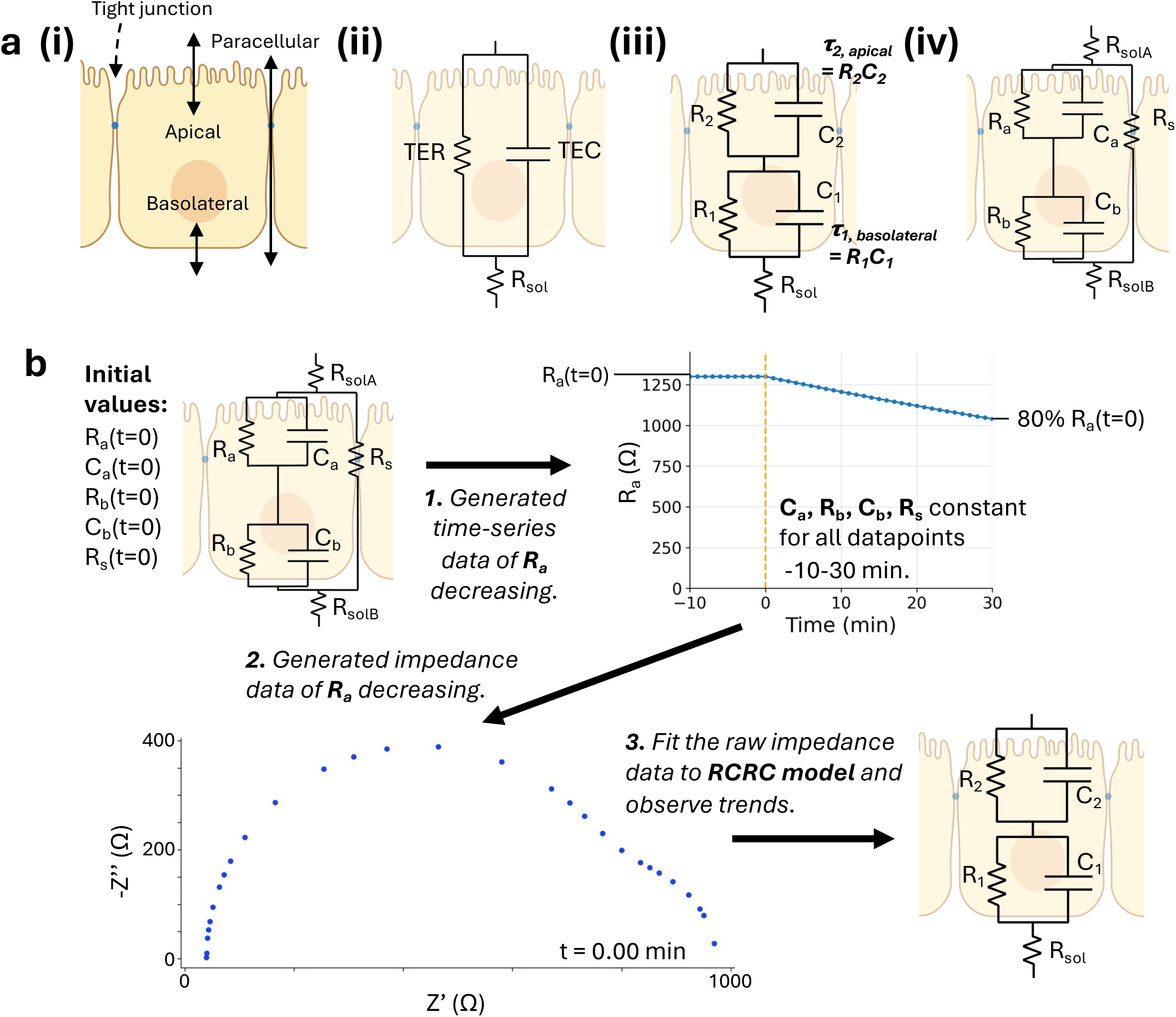
Method for determining how changes in apical resistance, R_a_, and basolateral resistance, R_b_, affect lumped parameters in RCRC model. (a) Apical, basolateral, and paracellular transport pathways across epithelia (i) have been modeled using various resistor–capacitor circuits proposed by us and others (ii–iv) [20–23]. We use the 7P model (iv) to gain insight into how the RCRC model (iii) reveals apical and basolateral electrophysiology in 16HBE cells. Schematics created partially with https://Biorender.com and adapted from [23]. (b) The method follows a three step process: (1) Generated time series data of R_a_ exponentially declining, (2) Generated impedance spectra for each time point, (3) Fit the raw impedance spectra to the RCRC model. The same process was also repeated for R_b_.

Because epithelial diseases frequently arise from disruptions to ion transport pathways, they can be studied through changes in the electrical properties of the tissue. Epithelial electrophysiology has therefore long provided insight into the regulation and physiology of barrier tissues. Single-point transepithelial resistance (TER/TEER) measurements provide an instantaneous snapshot of overall barrier integrity, capturing gross conductive changes across the tissue at a single moment. Short circuit current or voltage measurements in an Ussing chamber extend this approach by tracking changes over minutes to hours, enabling sequential pharmacological perturbations [6]. More recently, electrochemical impedance spectroscopy (EIS) techniques which send a multi-frequency signal enable measurement of transepithelial resistance (TER) and capacitance (TEC) over several days between different epithelial samples (Fig. 1a, ii). Despite differences in acquisition strategy and temporal sampling, these approaches are limited to bulk measurements across the entire epithelial tissue and do not distinguish apical and basolateral electrical contributions. To distinguish these contributions, intracellular sharp electrodes are required—an approach that requires great skill and years of training—such that intracellular electrophysiology of epithelia has remained limited to isolated studies from a small number of laboratories working on carefully selected cells and drugs [7–21].

Work in iPSC-derived retinal pigment epithelium has demonstrated that apical, basolateral, and paracellular electrical impedances can be resolved when extracellular impedance spectroscopy is paired with intracellular voltage recordings using the 7P-EIS technique [22]. However, this system requires complex electrophysiology hardware, tight intracellular access, and several hours of preparation and measurement for each sample. These limitations have prevented broader adoption and continue to restrict scalability, despite the well-established asymmetry of apical and basolateral channel and transporter expression in polarized epithelia and the added interpretability afforded by membrane-resolved measurements. To overcome these constraints in measurement speed and biological interpretability, we developed extracellular electro-chemical impedance spectroscopy (EEIS), which uses extracellular EIS coupled with fitting to a two-resistor, two-capacitor (RCRC) circuit to extract apical and basolateral membrane permeabilities. EEIS provides membrane-specific resolution from a single extracellular spectrum and quantifies reproducible permeability changes within 30 minutes, solving an important limitation with existing epithelial electrophysiology. We demonstrate that EEIS performs robustly across both simulated and biological experiments in wildtype 16HBE bronchiolar epithelial monolayers, enabling apical and basolateral membrane dynamics to be resolved through two electrical time constants, ***τ***_2_ and ***τ***_1_, respectively. Under conditions of CFTR activation and blocking with forskolin (10 µM) and GlyH-101 (10 µM), respectively, as well as during basolateral pore formation with nystatin (10 µM), membrane-specific perturbations are conserved cleanly in these time constants, while the assumption of negligible paracellular permeability shifts remains valid. Finally, we show that EEIS extends naturally to therapeutic evaluation, enabling rapid, ***τ***-based, membrane-resolved screening of pharmacological interventions for cystic fibrosis.

## 2 Results

### 2.1 Computational Simulation

Apical, basolateral, and paracellular transport pathways across epithelia (Fig. 1a, i) have long been represented using equivalent resistor–capacitor circuit models (Fig. 1a, ii–iv), which provide a framework for interpreting impedance spectra in terms of underlying membrane and barrier properties [20–23]. Among these, the seven-parameter (7P) epithelial model (Fig. 1a, iv) has been shown to correlate apical and basolateral membrane perturbations with distinct electrical elements, but requires intracellular voltage recordings to be fully resolved [22, 24]. In contrast, the reduced two-resistor, two-capacitor (RCRC) model (Fig. 1a, iii) can be fit using purely extracellular impedance measurements, but it remains unclear whether membrane-specific information is preserved following this parameter reduction.

To determine whether apical- and basolateral-specific contributions can still be resolved using a mathematical model with only 5 parameters instead of 7, we first compared the RCRC model in silico with the seven-parameter (7P) epithelial circuit model (Fig. 1a, iv). The 7P model requires intracellular voltage measurement to fully resolve, but converges to the RCRC model as the resistor corresponding to the para-cellular pathway, R_s_, approaches infinity, corresponding to no transport through the paracellular pathway and only through the apical and basolateral pathways. When R_s_ is finite, current flows through R_s_ and its impedance is combined with the apical and basolateral membrane impedances, absorbed into the effective RCRC parameters (*R*_1_*, R*_2_*, C*_1_*, C*_2_) during model reduction.

Because no closed-form analytical mapping exists between the parameters of the 7P and RCRC models under finite R_s_, it is not possible to explicitly relate *R_a_* and *R_b_* to individual RCRC elements (*R*_1_*, R*_2_*, C*_1_*, C*_2_*, R_sol_*). We hypothesized that dominant apical and basolateral membrane perturbations would remain separately identifiable in the reduced RCRC representation. We assessed this relationship empirically, decreasing the apical and basolateral membrane resistances in the 7P model, converting the data into impedance values, and then re-fitting to the RCRC model.

We generated physiologically relevant 7P measurements using a gridsearch of best fit solutions within ranges reported across epithelial systems (Methods section ‘Modeling EEIS’; Appendix A.1 & A.2). The resulting 7P measurements were then used as the initial values for a pseudo-experiment designed to test how imposed 20% apical or basolateral resistance decreases propagate into the RCRC measurements. R_sol_ accounts for series resistance from the extracellular media and produces only a rigid rightward offset in Nyquist space. Because this term does not participate in an RC element and therefore does not influence the extracted time constants, it was excluded from further analysis.

To validate the stability of the RCRC fitting procedure, we first simulated 10 impedance spectra with fixed 7P measurements (-10–0 min in Fig. 1b) and confirmed that the extracted RCRC parameters (*R*_1_*, R*_2_*, C*_1_*, C*_2_) and their derived time constants remained unchanged. We then introduced apical or basolateral perturbations by decreasing either *R_a_* or *R_b_* exponentially by 20% over 30 minutes of simulation (0– 30 min in Fig. 1b), with impedance sweeps analyzed sequentially in time to mimic an in vitro EEIS experiment as described in Fig. 1b. We chose 20% resistance steps to ensure detectable trends at the lower limit of resistance changes observed in literature [21, 22].

Impedance spectra generated by selective changes in *R_a_* or *R_b_* in the 7P model were subsequently fit to the reduced RCRC model using nonlinear least-squares optimization. Across all valid 7P solutions, the fitted RCRC parameters exhibited a consistent and conserved mapping: decreases in basolateral resistance were reflected as decreases in the larger time constant, ***τ***_1_, whereas decreases in apical resistance were reflected predominantly as decreases in the smaller time constant, ***τ***_2_. This mapping is consistent with the expected capacitance asymmetry: because tight junction proteins partition the cell closer to the apical surface, the basolateral membrane presents a larger area and therefore a larger capacitance (*C*_1_ *> C*_2_), which dominates the separation between ***τ***_1_ and ***τ***_2_.

Specifically, decreasing *R_b_* elicited a significant 11.6% reduction in ***τ***_1_ compared to less than 1% change in ***τ***_2_ (*p <* 0.001), whereas selective decreases in *R_a_* produced an 8.3% reduction in ***τ***_2_ and only a minor (3.6%) decrease in ***τ***_1_ over 30 min (*p <* 0.001; Fig. 2a, b). For both perturbations, a two-way repeated-measures ANOVA confirmed a significant ***τ*** *→* time-interval interaction (*p <* 0.001). Post-hoc paired *t*-tests (Holm–Bonferroni corrected) showing ***τ***_1_ significantly different from ***τ***_2_ at all intervals (*p <* 0.001), though the mean difference remained below 2% in the control intervals and grew progressively larger across subsequent time intervals. These differential responses persisted despite appreciable changes in the individual fitted resistive and capacitive parameters (Fig. A3). Individual resistive parameters do not map exclusively to isolated apical and basolateral resistance changes because perturbations to a single membrane domain induce coupled, nonlinear changes across fitted R and C values to achieve a minimum-error impedance fit. This behavior is an expected consequence of model reduction and motivates our focus on the composite time constants, which preserve membrane-specific dynamics. For resistance increases in *R_a_* and *R_b_*, we also observed conserved mapping: increases in basolateral resistance were reflected as increases in the larger time constant, ***τ***_1_, whereas increases in apical resistance were reflected as increases in the smaller time constant, ***τ***_2_ (Appendix A.4). Because this mapping held across the full ensemble of physiologically admissible 7P parameter sets, it provides a strong prediction for how membrane-specific perturbations should manifest experimentally, even in the absence of direct 7P measurements in 16HBE monolayers.

**Figure 2:**
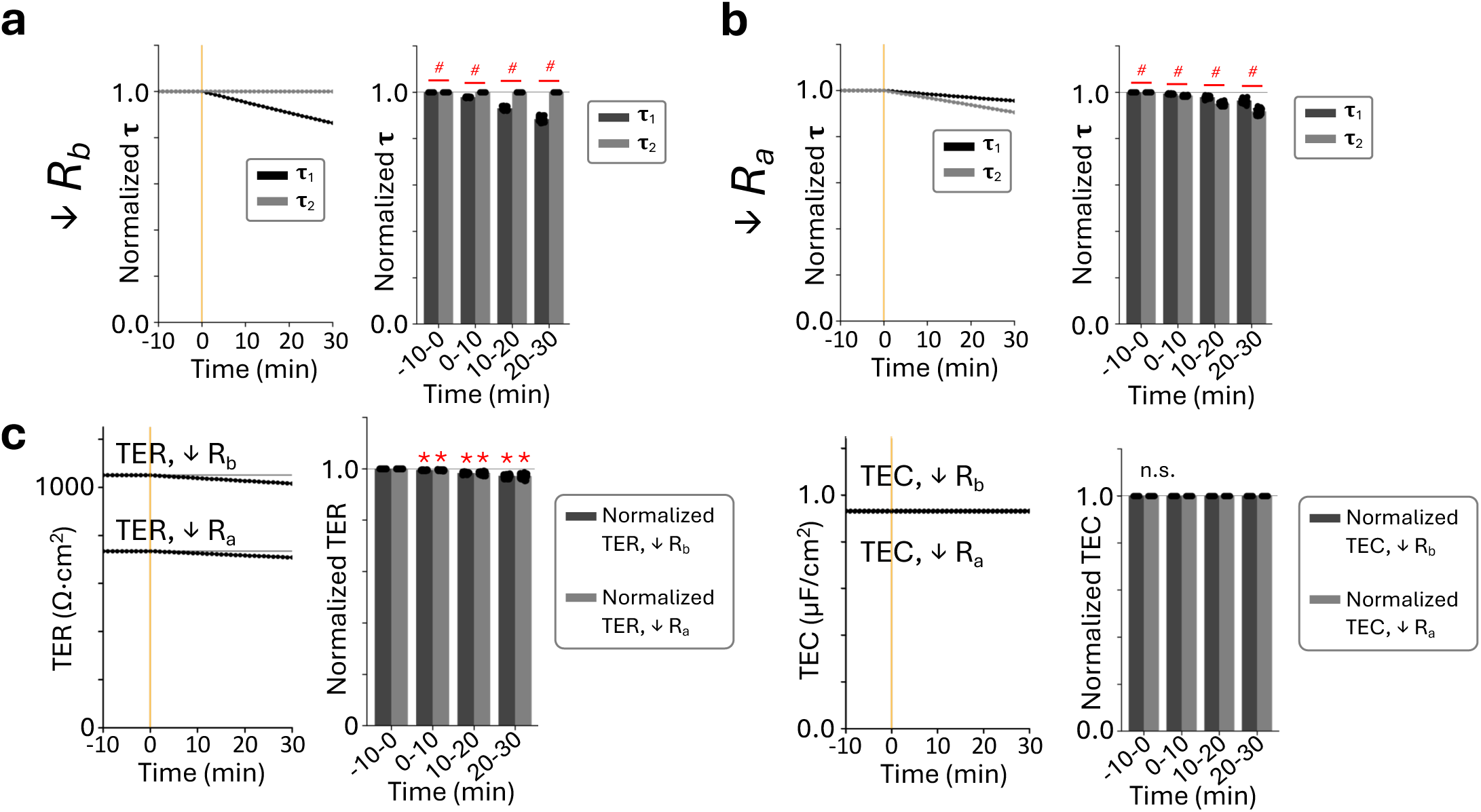
Effects of reducing apical and basolateral membrane resistances in simulation (n=58/group). (a) R_b_ decrease resulted in an 11.6% decrease in **τ**_1_ with <1% changes in **τ**_2_. (b) R_a_ decrease resulted in an 8.3% decrease in **τ**_2_ and a smaller 3.6% reduction in **τ**_1_ over the 30 min, despite significant changes in R_1_ and C_1_ (Appendix Fig. A3). For both (a) and (b), a two-way repeated measures ANOVA confirmed a significant tau × window interaction (p<0.001), with post-hoc paired t-tests (Holm-Bonferroni corrected) showing **τ**_1_ significantly exceeded **τ**_2_ at all intervals. ^#^p<0.001. (c) Representative transepithelial resistance (TER) measurements and bar charts of the average value within each 10 min interval showed significant decrease over time (one-way repeated measures ANOVA, p<0.001). Post-hoc paired t-tests with Holm-Bonferroni correction confirmed that all post-treatment intervals were significantly lower than the baseline interval (-10 to 0 min). *p<0.001. Representative transepithelial capacitance (TEC) measurements and bar charts showed no change in TEC (n.s.) Therefore, **τ**_1_ showed correlation with basolateral membrane changes and **τ**_2_ with apical changes.

Consistent with these membrane-resolved findings, simultaneous 20% reductions in both *R_a_* and *R_b_* produced significant decreases in transepithelial resistance (TER; one-way repeated measures ANOVA, *p <* 0.001; post-hoc paired *t*-tests with Holm– Bonferroni correction, *p <* 0.001 for all intervals 0–30 min relative to the *↑*10 to 0 min baseline; Fig. 2c). Transepithelial capacitance (TEC) showed no significant change over the same interval. This bulk response aligns with established understanding of TER, where permeabilizing the apical or basolateral membrane is observed as a net TER decrease.

Although transepithelial capacitance (TEC) can be measured robustly using impedance-based techniques, its physiological interpretation in polarized epithelia remains comparatively underexplored. Previous studies have suggested TEC as a possible indicator of cell surface area [25–29]. We report TEC here to note any unexpected changes, but focus biological interpretation on the TER and membrane-resolved time constants ***τ***_1_ and ***τ***_2_.

Nyquist plots generated after a 20% decrease in either *R_a_* or *R_b_* showed distinct qualitative shifts (Fig. 3). Because the model contains two RC time constants, where ***τ***_1_ is defined to be the larger time constant, the Nyquist response consists of two super-imposed semicircles. Selective reduction of *R_b_* predominantly altered the size of the right semicircle associated with ***τ***_1_ (Fig. 3a), whereas reduction of *R_a_* primarily modified the size of the left semicircle associated with ***τ***_2_ (Fig. 3b). Although points in the right semicircle shifted left when *R_a_* decreased (Fig. 3b), this is due to the *τ*_2_ semicircle shrinking, and the peak frequency of the right semicircle remained unchanged, conserving the underlying time constant. Therefore, each perturbation can be observed as one of the two time constants changing.

**Figure 3:**
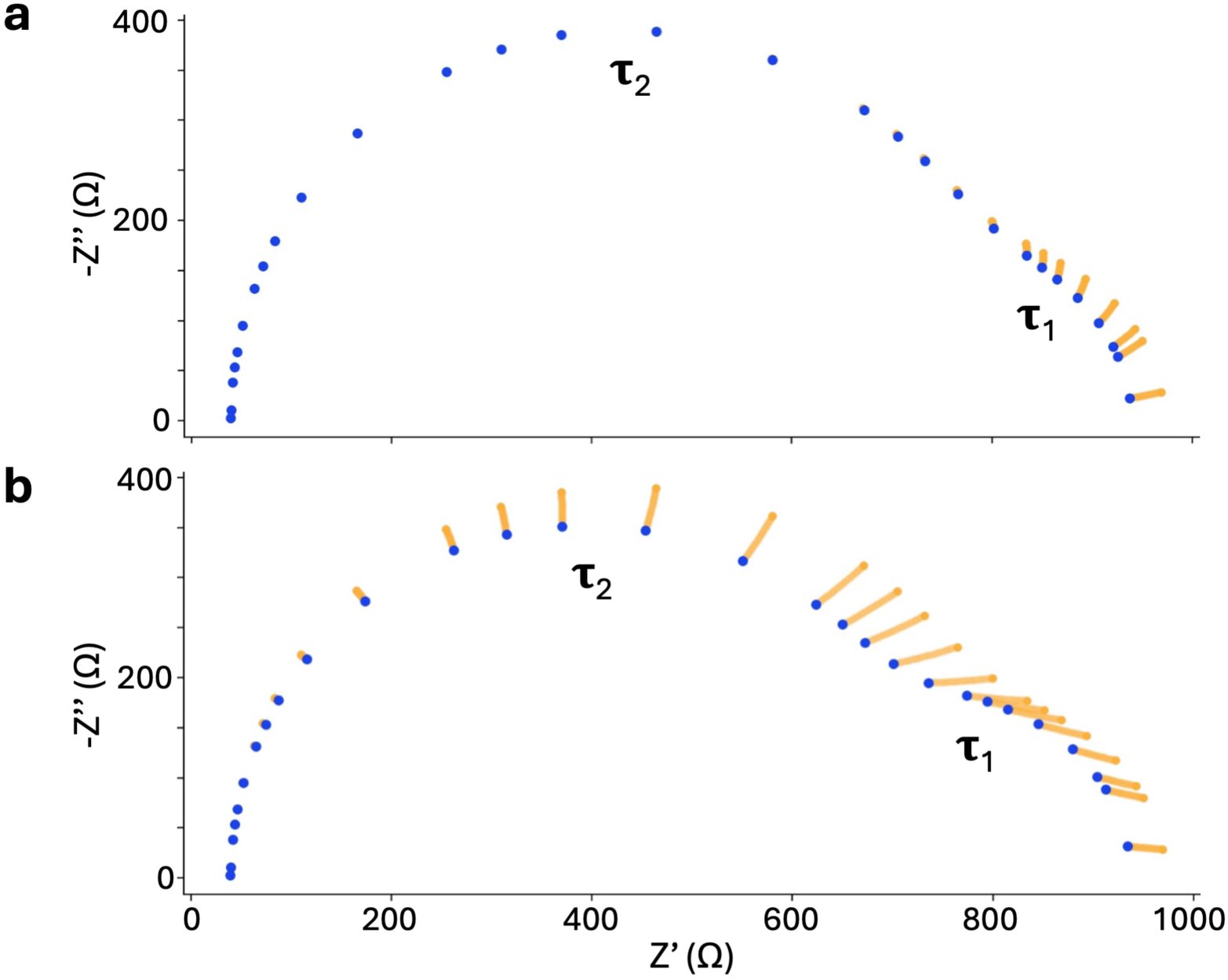
Representative Nyquist plot changes following reductions in apical and basolateral resistance in simulation. Negative imaginary impedance (-Z’’) vs. real impedance (Z’)). (a) R_b_ reduced to 80% of its initial value. (b) R_a_ reduced to 80% of its initial value. Nyquist plots shown are from the same starting impedances. Blue points denote the Nyquist response after the resistance reduction was complete, orange trace shows the trajectory of each point over the duration of the simulated experiment. Because the impedance spectrum contains two RC time constants, the response consisted of two semicircles: the left semicircle corresponds to the smaller time constant, **τ**_2_, while the right semicircle corresponds to the larger time constant, **τ**_1_. Selective reduction of R_b_ predominantly altered the right semicircle, whereas reducing R_a_ primarily reduced the left semicircle thus causing the right semicircle to shift toward lower Z’ values. Minor secondary effects on the **τ**_1_ semicircle observed do not affect the frequency corresponding to the apex of the semicircle, therefore leaving **τ**_1_ conserved.

Thus, in in silico experiments, TER behaved as expected for bulk epithelial metrics, and the RCRC-derived time constants preserve dominant apical and basolateral signatures that are not accessible through conventional readouts. These in silico results provided a principled basis for using these time constants as membrane-resolved indicators of permeability in subsequent in vitro experiments.

### 2.2 Increasing Apical Membrane Resistance in 16HBE

Based on our in silico results showing that apical perturbations primarily alter the smaller time constant *τ*_2_, we next examined the effects of blocking the apical CFTR channel in vitro, using 16HBE airway epithelial monolayers. To measure the maximum response, we first activated the CFTR channels with 10 µM forskolin using the experimental design in Fig. 4a. This caused a decrease in TER due to the decrease in resistance to chloride ions passing through the apical membrane. After the cells stabilized for 30 min, apical application of 5 µM or 10 µM GlyH-101 produced the expected increase in TER observed by others (Fig. 4b, i-ii) [30]. A 3 *→* 4 mixed ANOVA (between-subjects: treatment group [control/5 µM/10 µM]; within-subjects: time interval) revealed significant main effects of group, interval, and a significant group *→* interval interaction (all *p <* 0.001). Post-hoc pairwise t-tests with Holm-Bonferroni correction confirmed no baseline differences between any groups (-10-0 min, all *p >* 0.12). TER increased in a dose-dependent manner, with a greater response in the higher dose overall (*p* = 0.022). Notably, these TER trends matched the expected increase in resistance from blocking the CFTR channels, but were slower than short-circuit current responses typically observed in Ussing chambers [31], likely due to reduced convective mixing in the custom EEIS chambers [32].

**Figure 4:**
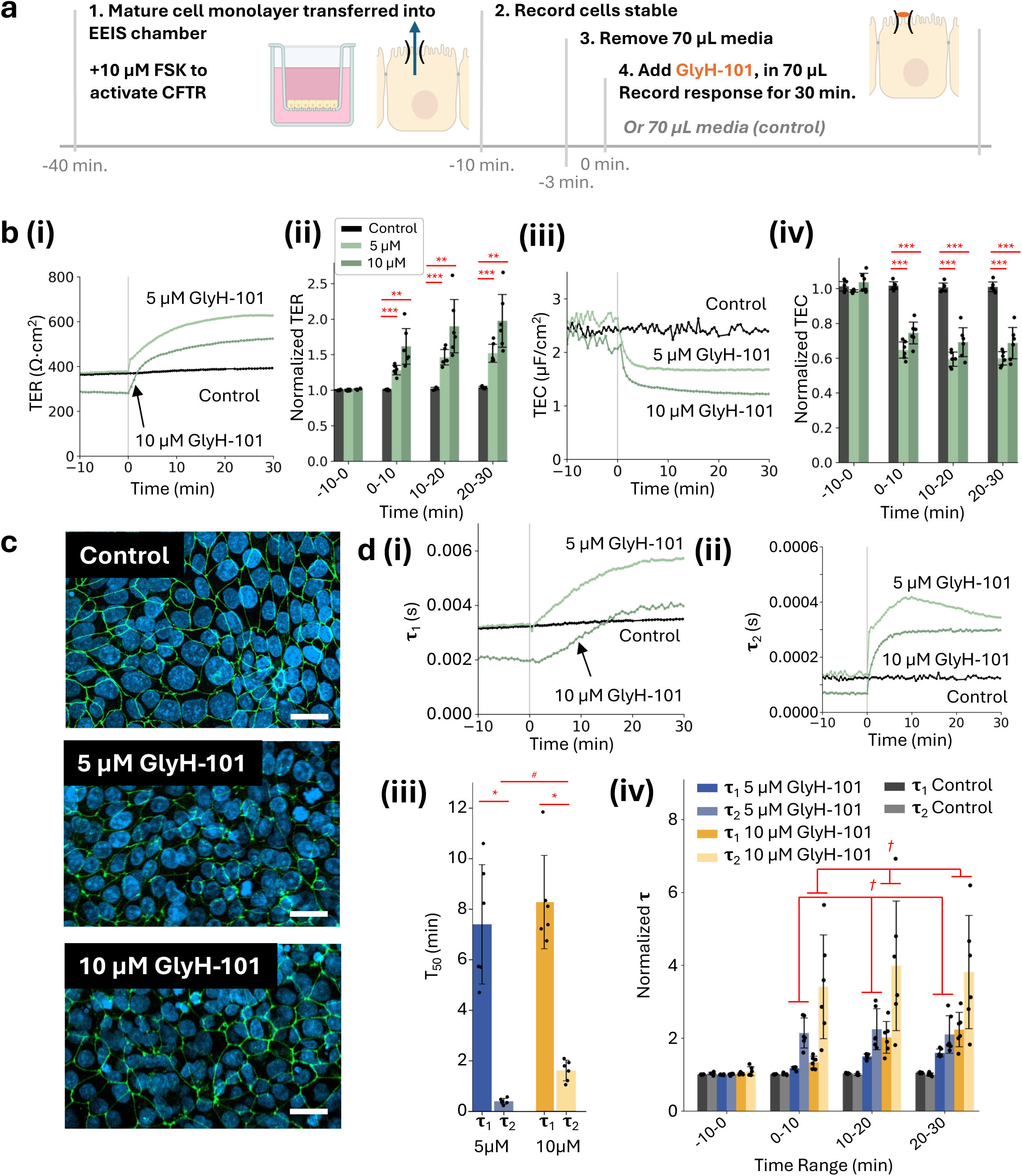
Effect of increasing apical membrane resistance with GlyH-101 on wild-type 16HBE (n=6/group). (a) Schematic illustrating the sequence and timing of forskolin and GlyH-101 perturbations and impedance acquisition during the experiment. (b, i-ii) 5 µM and 10 µM GlyH-101 resulted in a significant dose-dependent increase in TER in all post-treatment intervals. A 3×4 mixed ANOVA (between-subjects: treatment group [control/5 µM/10 µM]; within-subjects: time interval) revealed significant main effects of group (p<0.001), time interval (p_GG_<0.001), and a significant group × interval interaction (p<0.001). The 10 µM dose produced a significantly greater response than 5 µM overall (p=0.022). Post-hoc pairwise t-tests with Holm-Bonferroni correction confirmed no baseline differences between any groups (-10-0 min, all p>0.12). (b, iii-iv) 5 µM and 10 µM GlyH-101 resulted in a significant decrease in TEC at all post-treatment intervals, with a significant dose-dependent response overall (p=0.008). A 3×4 mixed ANOVA revealed significant main effects of group (p<0.001), time interval (p_GG_<0.001), and a significant group × interval interaction (p<0.001). Post-hoc pairwise t-tests with Holm-Bonferroni correction confirmed no baseline differences between groups (-10–0 min, all p>0.20). **p<0.01, ***p<0.001. (c) Following experiments, representative maximum intensity Z-projection confocal images of samples, showed intact tight junctions for all groups after the experiment, nuclei (DAPI, blue), Z0-1 (green), scale bar 20 µm. (d, i-ii) Representative measurements of **τ**_1_ and **τ**_2_ unnormalized. (d, iii) The half-maximal time, T_50_ of **τ**_1_ and **τ**_2_ revealed **τ**_2_ has a statistically faster T_50_ than **τ**_1_ for 5 µM and 10 µM doses (Paired t-test with Holm-Bonferroni correction, * p<0.05). The **τ**_2_ T_50_ of the 10 µM group was also statistically lower than the 5 µM group. t-test, ^#^p<0.05. (d, iv) Normalized **τ** values revealed GlyH-101 differentially affected **τ**_1_ and **τ**_2_ in a dose-dependent manner. A full factorial ANOVA (**τ** × group × interval) revealed a significant **τ** × group interaction (p=0.002), but no significant three-way interaction (p=0.980), indicating that the differential response of **τ**_1_ and **τ**_2_ to GlyH-101 was consistent across time intervals; post-hoc comparisons were therefore collapsed across post-treatment intervals. Both dose groups resulted in **τ**_2_ substantially exceeding **τ**_1_ increases (p<0.028), while in control samples, **τ**_1_ and **τ**_2_ remained comparable (n.s.) ^†^ p<0.05.

Across the same intervals, TEC exhibited rapid decreases *>* 20% within the first 10 min. A 3*→*4 mixed ANOVA revealed significant main effects of group, time interval, and a significant group *→* interval interaction (all *p <* 0.001). Post-hoc comparisons confirmed no baseline differences (-10-0 min, all *p >* 0.20). TEC stabilized within 5 minutes and remained lower than controls throughout the experiment (Fig. 4b, iii-iv). Representative immunofluorescence images of samples stained for the tight-junction protein ZO-1 (green) and nuclei (DAPI, blue) (Fig. 4c) demonstrated intact tight-junction organization, with no observable morphological damage following GlyH-101 exposure (i.e., at time *>* 30 min).

Representative unnormalized traces for *τ*_1_ and *τ*_2_ are shown in Fig. 4d, i-ii, highlighting the order-of-magnitude difference between the two time constants that allowed distinguishing independent *τ*_1_ and *τ*_2_ contributions, with *τ*_1_ consistently larger than *τ*_2_ across all groups. To compare the dynamics of the 5 µM and 10 µM responses, we calculated the half-maximal time (*T*_50_) for each trace (Fig. 4d, iii). For both GlyH-101 doses, *τ*_2_ reached its half-maximal response significantly faster than *τ*_1_, with mean *T*_50_ values 7 min (5 µM, *p* = 0.025) and 6.7 min (10 µM, *p* = 0.013) earlier than those of *τ*_1_. The 10 µM GlyH-101 group also exhibited a significantly slower *τ*_2_ T_50_ than the 5 µM group (*p* = 0.017), a pattern not seen in *τ*_1_, further supporting the assignment of *τ*_2_ to apical membrane dynamics (Fig. 4d, iii).

*τ*_1_ and *τ*_2_ normalized to their initial values at t=0 min, revealed GlyH-101 differentially affected *τ*_1_ and *τ*_2_ (Fig. 4d, iv). A full factorial ANOVA (*τ →* group *→* interval) revealed a significant *τ →* group interaction (p=0.002), but no significant three-way interaction (p=0.980), indicating that the differential response of *τ*_1_ and *τ*_2_ to GlyH-101 was consistent across time intervals; post-hoc comparisons were therefore collapsed across post-treatment intervals. Both dose groups resulted in *τ*_2_ substantially exceeding *τ*_1_ increases (*p <* 0.028), while in control samples, *τ*_1_ and *τ*_2_ remained comparable (n.s.) *^†^ p <* 0.05. Together, these analyses demonstrate the GlyH-101 induces both faster and larger magnitude responses in *τ*_2_, consistent with the membrane-specific assignments established in our in silico experiments.

To determine if we observed dose-dependent effects on *τ*_1_ and *τ*_2_ between the 5 and 10 µM dose, we performed linear regression of normalized *τ*_1_ and *τ*_2_ against GlyH-101 concentration at each time interval and tested for differences in dose sensitivity using a dose *→ τ* interaction term (Fig. 5). During the baseline interval (-10-0 min), both *τ*_1_ and *τ*_2_ remained near one across doses, and no significant difference in dose dependence between *τ*_1_ and *τ*_2_ was detected (dose x *τ*-type interaction, *p* = 0.114). In contrast, immediately following treatment (0–10 min), *τ*_2_ increased sharply with dose whereas *τ*_1_ showed only a modest dose-dependent increase, reflected in a highly significant dose x *τ*-type interaction (*p <* 0.001). This differential dose dependence persisted in all post-treatment intervals (all *p <* 0.001), indicating a significantly steeper dose–response relationship in *τ*_2_ vs. *τ*_1_. Together, these analyses demonstrate that GlyH-101 induces both faster and larger-magnitude dose-dependent responses in *τ*_2_, consistent with the membrane-specific assignments established in our in silico experiments.

**Figure 5:**
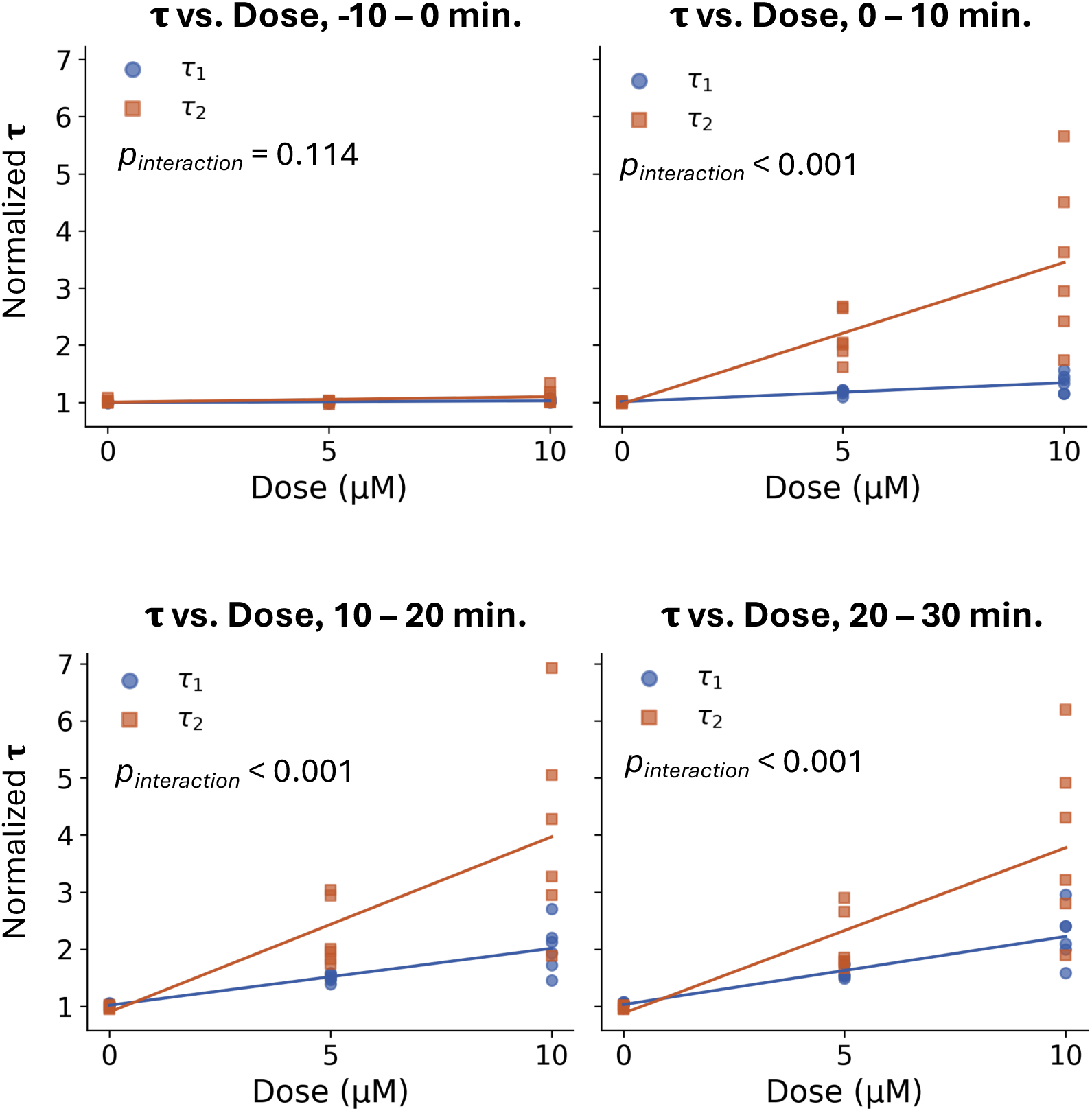
Effect of GlyH-101 dose on τ_1_ and τ_2_ in wild-type 16HBE cells (n = 6 per group). Dose–response relationships for **τ**_1_ (blue circles) and **τ**_2_ (orange squares) are shown across successive time intervals relative to dose (-10-0, 0-10, 10-20, and 20-30 min). Individual datapoints represent measurements from independent biological replicates, with solid lines indicating linear fits of **τ** versus GlyH-101 concentration at each interval. At baseline (-10-0 min), **τ**_1_ and **τ**_2_ exhibited comparable values and minimal dose dependence, with no significant difference in slopes (dose x τ interaction, p=0.114). Following treatment onset, **τ**_2_ displayed a significantly higher dose dependence than **τ**_1_ as reflected by significant dose × **τ** interaction terms (p<0.001 for all post-treatment intervals). These results indicate a time-dependent divergence in dose sensitivity between **τ**_1_ and **τ**_2_, with **τ**_2_ preferentially amplified at higher GlyH-101 concentrations.

### 2.3 Decreasing Basolateral Membrane Resistance in 16HBE

Because our in silico experiments indicated that basolateral perturbations would predominantly affect the larger time constant *τ*_1_, we next assessed impedance changes during basolateral permeabilization using 10 µM nystatin. Nystatin is known to insert into lipid membranes to form *↓*0.4 nm pores, thereby increasing basolateral ionic permeability without directly disrupting apical transport pathways [33]. Basolateral permeabilization with 10 µM nystatin, using the experimental design shown in Fig. 6a, produced the expected decrease in TER consistent with increased basolateral membrane permeability (Fig. 6b, i-ii). A 2 *→* 4 mixed ANOVA for TER revealed significant main effects of treatment group (control/nystatin), time interval, and a significant group × interval interaction (all *p <* 0.001). Post-hoc independent t-tests with Holm-Bonferroni correction confirmed no baseline difference (p=0.158), with nystatin-treated samples significantly lower than control at all post-treatment intervals (all *p <* 0.001, Fig. 6b, ii). These TER measurements validated the expected electrophysiological response to basolateral pore formation.

**Figure 6.**
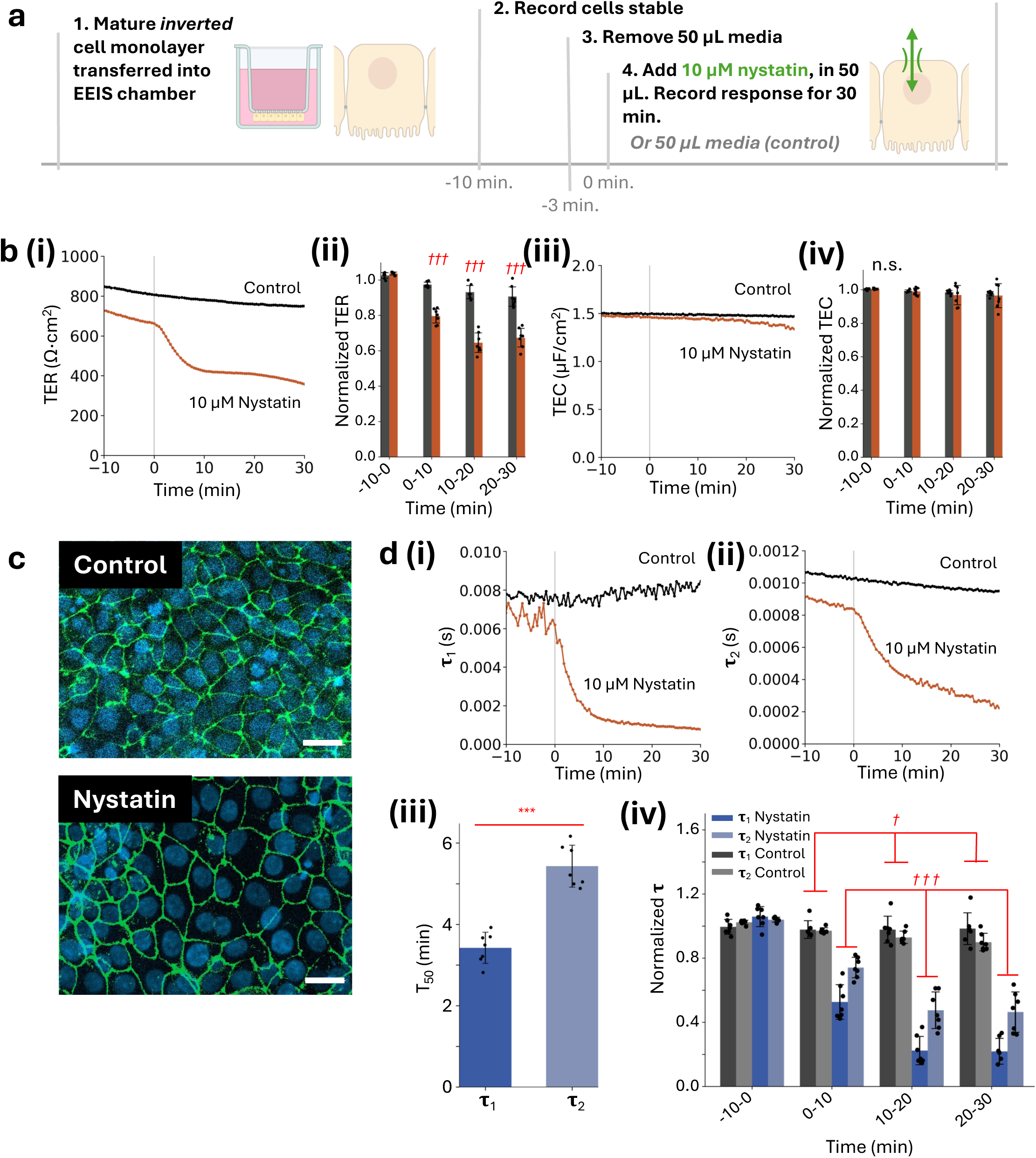
Effects of reducing basolateral membrane resistance with nystatin pores on wild-type 16HBE (n=7/group). (a) Schematic illustrating the sequence and timing of nystatin perturbations and impedance acquisition during the experiment. (b, i-ii) 10 µM nystatin resulted in a significant decrease in TER at all post-treatment intervals. A 2 × 4 mixed ANOVA for TER revealed significant main effects of group, time interval, and a significant group × interval interaction (all p<0.001). Post-hoc independent t-tests with Holm-Bonferroni correction confirmed no baseline difference (p=0.158), with nystatin-treated samples significantly lower than control at all post-treatment intervals (*^†††^* p<0.001). (b, iii-iv) TEC did not change significantly due to nystatin relative to the controls (2×4 mixed ANOVA group effect p=0.626; group × interval interaction p=0.789). (c) Following experiments, maximum intensity Z-projection confocal images of samples were taken. Representative images showed no change in morphology from staining of tight junction protein Z0-1 (green) and nuclei (DAPI, blue), scale bar 20 µm. (d, i-ii) Representative measurements of **τ**_1_ and **τ**_2_ unnormalized. (d, iii) The half-maximal time of the nystatin response, T_50_ of **τ**_1_ and **τ**_2_ revealed **τ**_1_ has a statistically smaller T_50_ than **τ**_2_. Paired t-test, *** p<0.001. (d, iv) Normalized **τ** values revealed nystatin differentially affected **τ**_1_ and **τ**_2_. A full factorial ANOVA (**τ** × group × window) revealed a significant **τ** × group interaction (p<0.001), but no significant three-way interaction (p=0.463), indicating that the differential response of **τ**_1_ and **τ**_2_ to GlyH-101 was consistent across time intervals; post-hoc comparisons were therefore collapsed across post-treatment intervals. In control samples, **τ**_1_ was modestly but significantly larger than **τ**_2_ (mean **τ**_1_=0.98 vs. **τ**_2_=0.93, p=0.042). In nystatin-treated samples, **τ**_2_ substantially exceeding **τ**_1_ (mean **τ**_1_=0.32 vs. **τ**_2_=0.56, p<0.001), consistent with a basolateral perturbation preferentially reducing **τ**_1_. Post-hoc paired t-tests with Holm–Bonferroni correction. ^†^ p<0.05, ^†††^ p<0.001.

Prior to addition of media or nystatin, no statistical differences were observed between control and nystatin groups; however, TER in the control condition exhibited a gradual *↓*15% decline over the 30 min experiment that was not observed in the apical channel blocking experiments. This slow drift may reflect increased sensitivity to media exchange in the inverted culture configuration. TEC did not change significantly due to nystatin relative to the controls (2 *→* 4 mixed ANOVA group effect *p* = 0.626; group *→* interval interaction *p* = 0.789, Fig. 6b, iii-iv).

Confocal images of representative 16HBE samples stained for ZO-1 (green) and nuclei (DAPI, blue) (Fig. 6c) showed intact tight-junction organization and no evidence of morphological damage following basolateral permeabilization (i.e., at *t >* 30 min). Together, these TER and TEC measurements serve as bulk electrophysiological validation of the expected response to basolateral perturbation.

Representative measurements of *τ*_1_ and *τ*_2_ (Fig. 6d, i-ii) demonstrate the characteristic order-of-magnitude separation between the two time constants, consistent with apical blocking experiments described previously, critical to distinguishing the two in fitting. Measurements of *τ*_1_ showed greater variability than those of *τ*_2_ prior to nystatin addition and in control samples, potentially attributable to the inverted culture configuration.

Normalized *τ* values revealed nystatin differentially affected *τ*_1_ and *τ*_2_. *τ*_1_ declined more rapidly, with a mean *T*_50_ occurring 2.00 min earlier than that of *τ*_2_ (*p <* 0.001; Fig. 6d, iii). A full factorial ANOVA (*τ →* group *→* interval) revealed a significant *τ →* group interaction (*p <* 0.001), but no significant three-way interaction (*p* = 0.463), indicating that the differential responses of *τ*_1_ and *τ*_2_ to nystatin were consistent across time intervals; post-hoc comparisons were therefore collapsed across post-treatment intervals. In nystatin-treated samples, *τ*_1_ decrease substantially exceeded *τ*_2_ (mean *τ*_1_=0.32 vs. *τ*_2_=0.56, *p <* 0.001), consistent with a basolateral perturbation preferentially reducing *τ*_1_. Thus, basolateral permeabilization with 10 µM nystatin elicited a significant basolateral-associated *τ*_1_ response compared with the apical-associated *τ*_2_. Unlike in the apical perturbation controls, where *τ*_1_ and *τ*_2_ remained comparable, basolateral controls exhibited a slight drift, with *τ*_1_ modestly but significantly exceeding *τ*_2_ (mean *τ*_1_=0.98 vs. *τ*_2_=0.93, p=0.042). We suspect similar to the TER drift that this is due to increased sensitivity in the inverted culture configuration.

### 2.4 Therapeutic Recovery of Apical Membrane Blocking in Cystic Fibrosis 16HBE

To test whether EEIS can detect restoration of apical CFTR function, we compared cystic fibrosis 16HBE (CF 16HBE) with and without chronic treatment with elexacaftor/tezacaftor/ivacaftor (ETI). Both groups were activated with 10 µM forskolin at *t* = *↑*40 min and blocked with 10 µM GlyH-101 at *t* = 0 min (Fig. 7a). Prior to GlyH-101 addition, ETI-treated epithelia exhibited significantly lower baseline resistances than untreated CF 16HBE (Fig. 7c), consistent with restored CFTR activation by forskolin (t-test, *p <* 0.001). We also noticed, as in the nystatin experiments, measurements of *τ*_1_—particularly in untreated CF 16HBE—exhibited greater variability, consistent with reduced epithelial stability in CF monolayers (Fig. 7d,i-ii).

**Figure 7.**
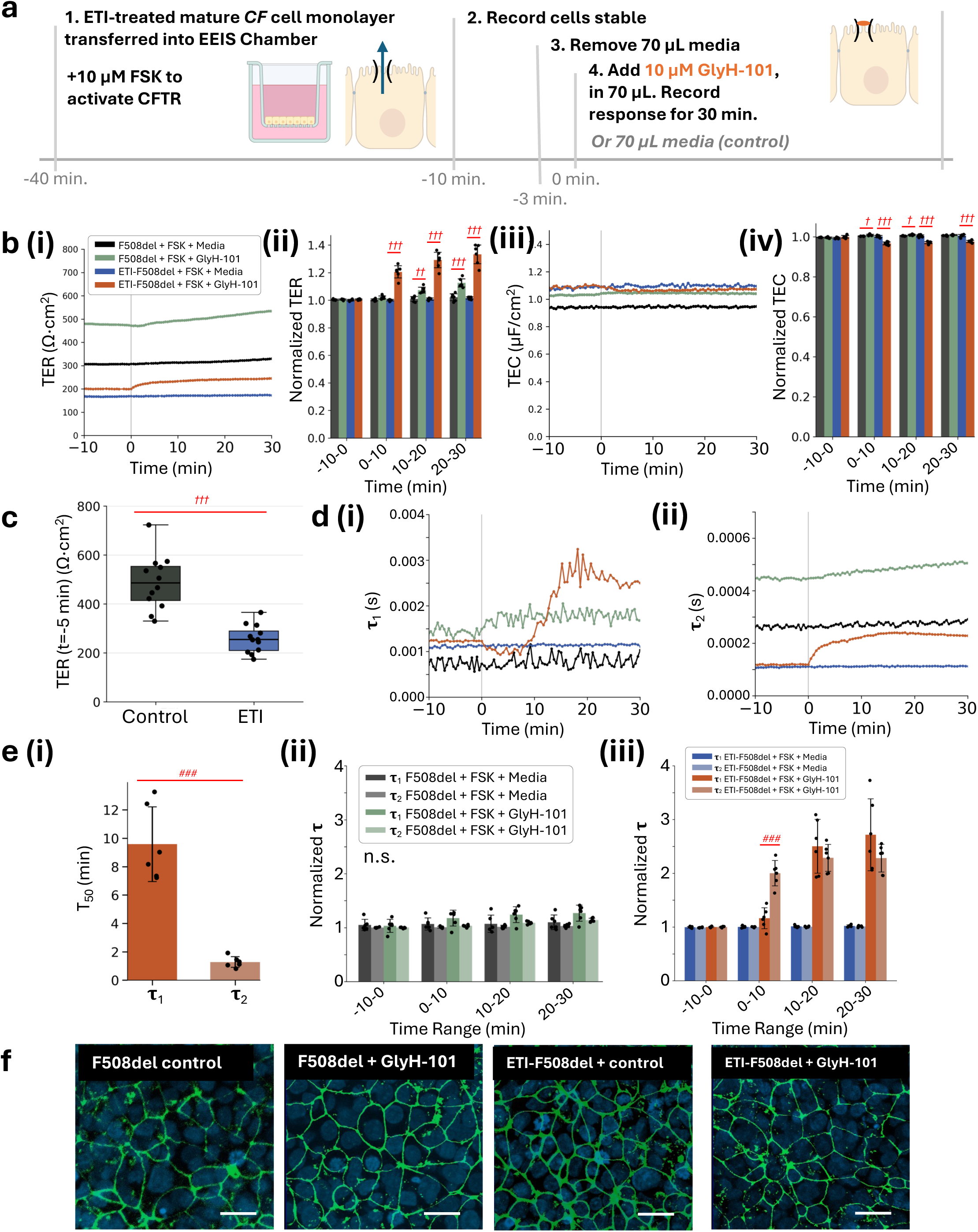
Effect of increasing apical membrane resistance with GlyH-101 with and without ETI treatment on CF 16HBE (n=6/group). (a) Schematic illustrating the sequence and timing of forskolin and GlyH-101 perturbations and impedance acquisition during the experiment. (b, i-ii) GlyH-101 TER response depends on ETI status. A 2×2×4 factorial ANOVA revealed a significant three-way group × ETI × window interaction (p<0.001). Decomposed 2×4 mixed ANOVAs showed significant group × interval interactions in both non-ETI and ETI-treated groups (both p<0.001). Post-hoc independent t-tests with Holm–Bonferroni correction confirmed no baseline differences in either group (all p>0.29). Following GlyH-101 addition, ETI-treated CF 16HBE exhibited a significant TER increase at all post-treatment intervals (all p<0.001). Non-ETI-treated CF cells also exhibited a TER increase, but with smaller magnitude and delayed onset, reaching significance at the 10–20 min (p=0.002) and 20–30 min intervals (p<0.001). ^††^p<0.01, ^†††^ p<0.001. (b, iii-iv) GlyH-101 significantly decreased TEC in ETI-treated cells and had a weaker effect in non-ETI cells. A 2×2×4 factorial ANOVA revealed a significant three-way group × ETI × window interaction (p<0.001), indicating that the GlyH-101 effect on TEC depends on ETI status. Decomposed 2×4 mixed ANOVAs revealed a significant group × interval interaction in both non-ETI (p<0.001) and ETI-treated groups (p<0.001). ETI-treated CF 16HBE showed a significant TEC decrease following GlyH-101 addition at all post-treatment intervals (all p<0.001). In non-ETI-treated CF cells, TEC showed a modest but significant increase at the 0–10 min (p=0.022) and 10–20 min intervals (p=0.022), but this effect did not persist at the 20–30 min interval (p=0.162). Post-hoc independent t-tests with Holm-Bonferroni correction confirmed no baseline differences in either group (all p>0.16). *^†^*p<0.05, *^†††^*p<0.001. (c) Stabilized TER values at -5 min, after 10 µM forskolin added at -40 min, had significantly lower resistances in the ETI-recovered groups prior to GlyH-101 addition, indicating the forskolin activation of the CFTR channel, causing a drop in resistance, was higher in the ETI treated groups (p<0.001). (d, i-ii) Representative measurements of **τ**_1_ and **τ**_2_ unnormalized. (e, i) The half-maximal time T_50_ of **τ**_1_ and **τ**_2_ of the ETI-treated GlyH-101 blocked group revealed **τ**_2_ has a statistically faster T_50_ than **τ**_1_. (t-test, ^###^ p<0.001). (e, ii) GlyH-101 differentially affected **τ**_1_ and **τ**_2_ depending on ETI status. A full factorial ANOVA (**τ** × group × ETI × window) revealed a significant four-way interaction, as well as significant **τ** × group × ETI (p<0.001) and **τ** × group interactions (both p<0.001). Decomposed factorial ANOVAs by ETI status showed no significant **τ**-related interactions in non-ETI-treated cells (**τ** × group, p=0.308; **τ** × group × window, p=0.950), indicating that **τ**_1_ and **τ**_2_ did not respond differentially to GlyH-101 in untreated CF cells. In ETI-treated cells, both **τ** × group (p<0.001) and **τ** × group × interval (p<0.001) interactions were significant, confirming a time-dependent differential response. Post-hoc paired t-tests with Holm–Bonferroni correction showed that GlyH-101 induced a significant and immediate response in ETI-treated CF 16HBE, with **τ**_2_ substantially exceeding **τ**_1_ at the first post-treatment interval (normalized **τ**_2_=2.01 vs. **τ**_1_=1.17, p<0.001), although this difference was transient as **τ**_1_ increased to match **τ**_2_ at later intervals (all p>0.50). In both control groups, **τ**_1_ and **τ**_2_ remained comparable across all intervals (all n.s.) ^###^p<0.001. (f) Following experiments, maximum intensity Z-projection confocal images of samples were taken. Representative images showed intact tight junctions for all groups, DAPI (blue), Z0-1 (green), scale bar 20 µm. Therefore, we observed recovery of CFTR function indicated by the faster **τ**_2_ T_50_ in the ETI-treated GlyH-101 blocked group over **τ**_1_ and the significantly larger response in the 0-10 min interval.

Representative TER and TEC measurements are shown in Fig. 7b,i and iii. A 2 *→* 2 *→* 4 factorial ANOVA revealed a significant three-way group *→* ETI *→* interval interaction (*p <* 0.001). Decomposed 2 *→* 4 mixed ANOVAs showed significant group *→* interval interactions in both non-ETI and ETI-treated groups (both *p <* 0.001). Post-hoc independent t-tests with Holm–Bonferroni correction confirmed no baseline differences in either group (all *p >* 0.29). Following GlyH-101 addition, ETI-treated CF 16HBE exhibited a significant TER increase at all post-treatment intervals (all *p <* 0.001). Non-ETI-treated CF cells also exhibited a TER increase, but with smaller magnitude and delayed onset, reaching significance at the 10–20 min (p=0.002) and 20–30 min intervals (*p <* 0.001).

TEC responses also distinguished the groups (Fig. 7b, iv). Both ETI and non-ETI conditions showed strong TER responses differing primarily in magnitude, while TEC discriminated the two conditions more clearly: the non-ETI group effect (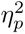 = 0.453) was substantially weaker than the ETI group effect (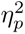 = 0.836). A 2 *→* 2 *→* 4 factorial ANOVA revealed a significant three-way group *→* ETI *→* interval interaction (*p <* 0.001), indicating that the GlyH-101 effect on TEC depends on ETI status. Decomposed 2 *→* 4 mixed ANOVAs revealed a significant group *→* interval interaction in both non-ETI (*p <* 0.001) and ETI-treated groups (*p <* 0.001). ETI-treated CF 16HBE showed a significant TEC decrease following GlyH-101 addition at all post-treatment intervals (all *p <* 0.001) with the Greenhouse–Geisser-corrected interval effect losing significance (*p_GG_* = 0.107), suggesting that the TEC response in ETI-treated cells was rapid and stable rather than progressive. In non-ETI-treated CF cells, TEC showed a modest but significant increase at the 0–10 min (p=0.022) and 10–20 min intervals (p=0.022), but this effect did not persist at the 20–30 min interval (p=0.162). Post-hoc independent t-tests with Holm-Bonferroni correction confirmed no baseline differences in either group (all *p >* 0.16).

GlyH-101 differentially affected *τ*_1_ and *τ*_2_ depending on ETI status (Fig. 7d-e). A full factorial ANOVA (*τ →* group *→* ETI *→* interval) revealed a significant fourway interaction, as well as significant *τ →* group *→* ETI (*p <* 0.001) and *τ →* group interactions (both *p <* 0.001). Decomposed factorial ANOVAs by ETI status showed no significant *τ*-related interactions in non-ETI-treated cells (*τ →* group, p=0.308; *τ →* group *→* interval, p=0.950), indicating that *τ*_1_ and *τ*_2_ did not respond differentially to GlyH-101 in untreated CF cells. In ETI-treated cells, both *τ →* group (*p <* 0.001) and *τ →* group *→* interval (*p <* 0.001) interactions were significant, confirming a time-dependent differential response. This is consistent with the nearly 10 min delay observed in the *τ*_1_ increase (Fig. 7d, i), relative to the *τ*_2_ immediate increase. Post-hoc paired t-tests with Holm–Bonferroni correction showed that GlyH-101 induced a significant and immediate response in ETI-treated CF 16HBE, with *τ*_2_ substantially exceeding *τ*_1_ at the first post-treatment interval (normalized *τ*_2_=2.01 vs. *τ*_1_=1.17, *p <* 0.001), although this difference was transient as *τ*_1_ increased to match *τ*_2_ at later intervals (all *p >* 0.50). In both control groups, *τ*_1_ and *τ*_2_ remained comparable across all intervals (all n.s.)

Additionally, the half-maximal times for *τ*_1_ and *τ*_2_ in ETI-treated CF epithelia were statistically indistinguishable from those measured in wild-type cells undergoing the same forskolin and GlyH-101 protocol (Fig. 4d, iv), indicating recovery of the EEIS impedance phenotype associated with functional apical CFTR. Although, the *τ*_1_ delay observed in the ETI-treated CF group is not present in the wild-type response. Confocal images of representative samples stained for ZO-1 (green) and nuclei (DAPI, blue) (Fig. 7f) showed intact tight-junction organization and no clear evidence of morphological damage following apical permeabilization or ETI treatment (i.e., at *t >* 30 min), although differences in localization of ZO-1 in CF epithelia have been previously reported [31].

## 3 Discussion

EEIS consistently resolves apical and basolateral electrical behavior through *τ*_2_ and *τ*_1_, respectively. Apical perturbations, including CFTR blocking with GlyH-101, produced larger and faster changes in *τ*_2_, and basolateral permeabilization with nystatin produced larger and faster changes in *τ*_1_ as measured by *T*_50_ and normalized *τ*changes. These signatures were dose dependent in GlyH-101, reproducible across biological replicates, and robust to natural variation in initial resistance and capacitance. Together with our in silico analysis, these findings establish a clear mechanistic mapping between the RCRC model and the underlying epithelial membrane domains. As supplementary validation, we also examined the response to 10 µM forskolin alone, which increases both apical CFTR conductance and basolateral Na^+^–K^+^–2Cl*^→^*cotransporter activity. Under this condition—where both membrane permeabilities change simultaneously—*τ*_1_ and *τ*_2_ declined with similar kinetics (Fig. A5), but approached significance in the first 10 min. with faster and larger normalized *τ*_1_ responses. Although not central to the main results, this side experiment confirms that when membranes are modulated from an addition on the apical side, we can still observe basolateral changes, reinforcing our interpretation that the asymmetric responses observed under selective perturbations reflect true membrane-specific dynamics.

EEIS also uncovered an unexpected and previously unreported decrease in transepithelial capacitance (TEC) following CFTR blocking. TEC decreased with GlyH-101 addition, but did not change during basolateral pore formation (Fig. 4b, iii-iv and Fig. 7b, iii-iv). Because TEC captures the combined capacitive contribution of both membranes, this pattern suggests that apical CFTR blocking reduces the effective apical membrane capacitance, whereas basolateral permeabilization primarily alters conductance. One possible explanation is that CFTR blocking modifies the contribution to charge storage, whereas nystatin creates a predominantly resistive shunt that does not significantly alter capacitance.

Importantly, the therapeutic experiments demonstrate that EEIS can detect recovery of apical CFTR function. ETI-treated CF epithelia exhibited TER, TEC, and *τ*_2_ dynamics that matched wild-type (WT) 16HBE responses, including a rapid *τ*_2_ increase after apical CFTR blocking and significantly lower baseline resistances following forskolin activation (Fig. 7). The half-maximal times of *τ*_1_ and *τ*_2_ in ETI-treated CF cells were statistically indistinguishable from WT, indicating recovery of the impedance phenotype. These results highlight the utility of EEIS as a rapid, extracellular approach for quantifying pharmacological rescue of apical CFTR function without the need for intracellular access or complex electrophysiology.

Our analysis assumes that paracellular resistance remains relatively stable, an assumption supported by both simulations and empirical data under the perturbations used here. Nonetheless, it remains unknown how deliberate modulation of tight-junction permeability would influence the mapping between *τ*_1_, *τ*_2_, and the membrane domains. Addressing this question will require targeted junctional perturbations together with expanded circuit modeling. Additionally, while we focused on GlyH-101 and nystatin as representative apical and basolateral perturbations, broader validation using other modulators and epithelial types will be necessary to generalize this framework.

Overall, this work establishes EEIS as a rapid, scalable, and biologically interpretable method for resolving membrane-specific electrical dynamics in polarized epithelia. By combining in silico modeling with targeted experimental perturbations, we provide a foundation for applying this approach to therapeutic screening, disease modeling, and mechanistic studies of epithelial transport.

## 4 Materials and Methods

### 4.1 RCRC Model Fitting

Raw impedance spectra were exported from NOVA 2.1 (Metrohm, 2024) and fit to the circuit in Fig. 1a, iii using MATLAB code available at https://github.com/chien0507/ extracellularEIS/tree/main [23].

Briefly, the modeled impedance is

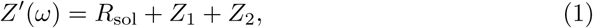

where

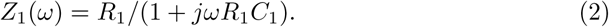

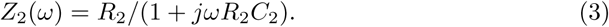

Model parameters were obtained by minimizing the nonlinear least-squares cost function

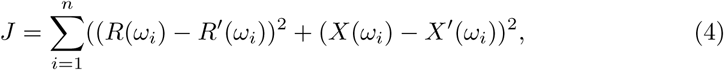

using MATLAB’s lsqcurvefit with the optimization settings in Table 1, modified from [22]. Across all biological experiment fits, the residual norm (resnorm) values were less than 0.1, with a median of 0.0194 and an interquartile range (IQR) of 0.0268 (n=5003); across simulated experiment fits, the residual norm (resnorm) values were low, with a median of 6.41E-17 and an interquartile range (IQR) of 1.06E-16 (n=9512). indicating stable and well-constrained parameter recovery under the selected bounds.

**Table 1:** Settings for nonlinear least squares fit. These non-default settings are configured in Matlab’s ‘optimoptions’ function and are applied to the ‘lsqcurvefit’ algorithm.

| Parameter | Value | Description |
| --- | --- | --- |
| MaxIterations | $3 \times 10^3$ | Maximum sets of parameter values ($R_s, R_1, R_2, C_1, C_2$) tried before reporting the best. |
| MaxFunctionEvaluations | $3 \times 10^3$ | Maximum number of function evaluations allowed. |
| FunctionTolerance | $3 \times 10^{-9}$ | Tolerance for convergence based on the function value. |
| OptimalityTolerance | $5 \times 10^{-7}$ | Tolerance for converging based on the gradient of the error function. |
| StepTolerance | $1 \times 10^{-9}$ | Tolerance for converging based on the step size of the error function. |

Fitting bounds were chosen to encompass previously reported epithelial membrane properties [22, 29, 34–39]:

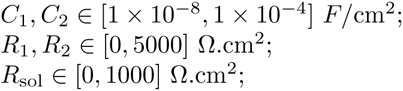

### 4.2 Modeling EEIS

To establish the relationship between the seven-parameter (7P) epithelial circuit model and the two-resistor, two-capacitor (RCRC) model, we first generated a database of physiologically relevant 7P parameter sets (see Fig. A1, A2). Parameter bounds were determined from previously published impedance measurements in mature 16HBE monolayers and from ranges defined in prior literature, as described in Methods section ‘RCRC Model Fitting’. A grid search across these bounds was used to identify all valid 7P solutions that reproduced the experimentally measured impedance spectra from four biological samples.

Using each of the 58 valid 7P solutions as an initial condition, we constructed a pseudo-experiment spanning 40 min. For the first 10 min, all 7P parameters were held constant. For the subsequent 30 min, either the apical resistance (*R_a_*) or basolateral resistance (*R_b_*) was reduced exponentially to 80% of its initial value to mimic a targeted change in membrane permeability. A 20% reduction was chosen because it exceeds the baseline drift typically observed in mature 16HBE monolayers while remaining small enough to test EEIS sensitivity to modest, membrane-specific changes. At each simulated minute, the corresponding 7P parameters were used to generate impedance spectra at the same frequencies used experimentally.

Each synthetic impedance spectrum was then fit to the RCRC model using previously published MATLAB routines [23], producing time-resolved estimates of R_1_, R_2_, C_1_, C_2_, the derived time constants *τ*_1_ = R_1_ *·* C_1_ and *τ*_2_ = R_2_ *·* C_2_, *TER* = *R*_1_ + *R*_2_, 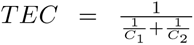. These simulations allowed us to map apical- and basolateral-specific perturbations in the 7P model onto changes in the RCRC-derived time constants.

### 4.3 16HBE Cell Culture for Apical Perturbation

16HBE14o-human bronchial epithelial cells expressing wild-type CFTR (16HBE-WT) (MilliporeSigma, SCC150) were seeded at 250,000 cells/well in 12 mm Transwell® with 0.4 µm pore PET membrane insert (Corning, 3460) and cultured following recommendations made by the CFFT [40], except flasks were not coated with collagen. Cells were grown in liquid-liquid conditions with media composed of 89% MEM media (Thermo Fisher Scientific, 11095098), 10% fetal bovine serum, premium, heat inactivated (R&D Systems, S11150H or Corning, 35-015-CV), and 1% penicillin-streptomycin (5,000 U/mL) (Thermo Fisher Scientific, 15070063) with media changes three times per week until cells were confluent [40]. Cultures were used for up to 12 weeks after thawing. Maturity was confirmed using the EndOhm (12 mm) (World Precision Instruments, EndOhm-12G) and EVOM2, measuring resistances over 500 Λ*·*cm^2^. At the start of the EEIS experiment, 10 µM forskolin was added to achieve maximal CFTR activation in 16HBE monolayers, as established in prior studies [31, 41]. After stabilization, GlyH-101 stock solution was added to the apical compartment to reach half-maximal or maximal blocking at 5 or 10 µM, consistent with previously studies [42].

### 4.4 16HBE Cell Culture for Basolateral Perturbation

16HBE14o-Human bronchial epithelial cells expressing wildtype CFTR (16HBE-WT) (MilliporeSigma, SCC150) were cultured under identical conditions to the apical-perturbation experiments, except they were seeded on the undersurface of 12-well cell culture inserts with 0.4 µm pores coated with collagen I (Corning, 3460) following the established inverted culture protocol of the McCarty laboratory[43] and forskolin was not added at the start of the EEIS experiment. During the EEIS experiment, nystatin was added to the basolateral-facing bath at a final concentration of 10 µM, selected based on a preliminary dose–response study (data not shown).

### 4.5 16HBE Cell Culture for ETI Experiments

16HBE14o-human bronchial epithelial cells expressing wildtype CFTR (16HBE-WT) (MilliporeSigma, SCC150) and 16HBE cells expressing F508del-CFTR on a V470 background (16HBE-CF) (CFF, CFF-16HBEge CFTR F508del/V470) were used for ETI experiments. Both cell types were cultured inside 12-well cell culture inserts with 0.4 µm pores following the same seeding and culture protocol as that for the apical perturbation experiments, except medium was supplemented beginning on day 6 with 5 µM elexacaftor and 18 µM tezacaftor, followed by 1 µM ivacaftor on day 7 for ETI groups, and control (non-ETI) groups received identical media without modulators [43].

### 4.6 EEIS on Biological Samples

Cell culture inserts were transferred into a custom electrophysiology chamber containing 3.5 mL room-temperature RPMI-1640 medium (Corning, 10-040-CV) in the basolateral compartment and 0.7 mL in the apical compartment. Platinum wire loops served as current-carrying electrodes, and platinum wires were used as voltage-sensing electrodes. A CAD rendering of the chamber is shown in Fig. A6. Previous work has shown consistent measurements and minimal error on a model electrical circuit and used to measure iPSC-derived retinal pigment epithelia and 16HBE cell monolayers with an EndOhm chamber [23], establishing mean absolute error and L2 norm of the residuals (resnorm) as ways to quantify error. EEIS measurements were started and the cells were then given 30 minutes to stabilize in the chamber, new media, and room temperature prior to experimentation, similar to established short-circuit current protocols [31].

After placement in the chamber, monolayers were allowed to stabilize for 30 min at room temperature, consistent with established short-circuit current protocols [31]. EEIS measurements were then collected for an additional 7 min to establish a stable baseline. At *t* = 0 min, a small volume of medium (*<* 100 *µ*L) was removed and replaced with an equal volume containing modulators or control vehicle. Control inserts confirmed that media exchange alone did not alter impedance. Samples were recorded for 30 min following addition.

EEIS measurements consist of electrochemical impedance spectroscopy sweeps and fitting to the two-resistor two-capacitor circuit described in Fig. 1a, iii. The impedance sweeps are from 2 Hz - 50 kHz at 4 µA amplitude with two frequencies per decade and 5 frequencies stacked at low frequencies (5sines setting) performed using a custom NOVA protocol [23]. The signal is sent and impedance is measured using the Multi Autolab cabinet with M204 multichannel PGSTAT and FRA32M modules modules (Metrohm, catalog number: AUT.MAC204.S, FRA32M.MAC.204.S).

### 4.7 Immunostaining and Confocal Microscopy

Immunostaining and confocal microscopy were performed as previously established [31, 43]. Cells seeded on cell culture inserts were fixed using 2% paraformaldehyde in 1x PBS for 10 min, followed by a 2-minute incubation with 1:1 methanol acetone. Samples were blocked using 2% BSA in PBS and incubated overnight at 4*^↓^*C with primary antibodies targeting ZO-1 (ThermoFisher, 33-9100). Samples were washed and incubated with secondary antibody Alexa Fluor-594 anti-mouse (1:500) for 1 hr (ThermoFisher, A11005). Transwells were cut and mounted with Vectashield antifade with DAPI (Vector Laboratories, H-1200-10). All steps were performed at room temperature. Images were collected using a laser scanning confocal microscope (Zeiss, LSM 900). Representative maximum Z projections (Max-Z) confocal images of 16HBE cells are shown.

### 4.8 T_50_ Calculation

For each sample, the time to half-maximal response (T_50_) was computed from the raw, non-windowed time-series data. The baseline value *y*_0_, was established as the value at *t* = 0. For increasing responses, the maximum value *A*_max_ was defined as the raw maximum of the signal over the 0–30 min interval. For decreasing responses, the minimum value *A*_min_ was defined as the raw minimum of the signal over this same time interval. T_50_ target value was defined as the midpoint between the baseline and the corresponding minimum or maximum:

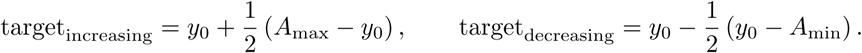

T_50_ was defined as the first post-baseline time point at which the measurements reached the 50% target value. Threshold crossings were identified by detecting adjacent samples that flanked the target (increasing for rising responses and decreasing for falling responses), and the crossing time was estimated by linear interpolation between these points. Bar plots display the mean value, with standard deviation as the error bars.

### 4.9 Statistical Analysis

All normalized values were computed relative to the steady-state measurement at *t* = 0 min, prior to modulator addition. Bar plots display the mean value within each time interval for each biological replicate, following conventions used for Ussing short-circuit current analyses, with error bars representing the standard deviation across replicates. All *n* = values reported throughout refer to biological replicates.

Statistical analyses were performed in Python using the SciPy, Statsmodels, and Pingouin packages, complete ANOVA results and post-hoc tests for in silico experiments are reported in Tables A3, A4. Complete ANOVA results and post-hoc tests for in vitro experiments are reported in Tables A7, A8.

## 5 Conclusion

This work establishes extracellular electrochemical impedance spectroscopy (EEIS) as a rapid, extracellular, and biologically interpretable approach for resolving apical and basolateral membrane electrophysiology in 16HBE via membrane-specific time constants, overcoming the lumped barrier readouts of conventional TEER and EIS measurements. In contrast to 7P-EIS recordings, which require invasive electrodes, extensive setup, and low-throughput measurement protocols, EEIS offers a scalable alternative capable of domain-specific resolution without intracellular access.

By combining electrical modeling, in silico validation, and targeted perturbations of CFTR channels and basolateral permeability, we show that *τ*_2_ reflects apical membrane changes while *τ*_1_ reflects basolateral electrophysiology in 16HBE cells. This membrane-specific resolution allows EEIS to quantify therapeutic responses, as demonstrated by the recovery of apical CFTR function in ETI-treated cystic fibrosis epithelia. Because EEIS operates extracellularly and supports high sample throughput, it may enable screening of CFTR-restoring therapies directly on patient-derived cultures, providing a scalable complement or alternative to Ussing chamber assays.

Looking forward, the ability of EEIS to extract membrane-specific electrical behavior from a single extracellular spectrum positions this technique for integration into organ-on-chip platforms, microfluidic culture systems, and longitudinal studies of epithelial barrier function. Such integration could enable rapid assessment of apical and basolateral physiology in engineered tissues, expanding the utility of impedance-based diagnostics and supporting the development of targeted therapeutics for epithelial diseases.

## Acknowledgements

Thank you to the Montgomery machine mall for providing equipment to mill the chamber, and facilities and staff supporting the Petit Institute for Bioengineering and Bioscience and Emory University.

## Declarations

### Data availability

Raw impedance data for in silico and in vitro experiments are available on Github here: https://github.com/chien0507/memResolvedEEIS.git

### Materials availability

Gibco FBS was discontinued midway through experimentation, and Corning FBS was used moving forward. Controls have established no statistical difference observed in the TER values of the samples.

### Code availability

Fitting code has been previously published and is available on Github [23].

### Author contribution

A.C. designed and performed experiments, analyzed data, and wrote the manuscript. G.C. assisted in inverted culture technique. E.L. machined the electrophysiology chamber. H.K. did error analysis and assisted manuscript editing. N.M. assisted in results interpretation and acquired funding. C.F. assisted in results interpretation, writing, and acquired funding.

## Appendix A Additional Figures

**A.1 Method for determining physiologically relevant 7P initial values.**

**A.2 Assessing Variability in Modeling Initial Values**

**A.3 Additional Modeling Results: *R*_1_, *R*_2_, *C*_1_, *C*_2_ response to decreasing *R*_a_ and *R*_b_**

**A.4 Additional Modeling Results: Increasing *R*_a_ and *R*_b_**

**A.5 Forskolin Results**

**Table A1:** Mixed ANOVA results for forskolin experiments (*n* = 6 per group). 2 *→* 4 mixed ANOVAs (between: group; within: interval) were used for TER, TEC, *τ*_1_, and *τ*_2_. A sepa- rate mixed ANOVA (between: group; within: *τ*type) was used for the *τ*_1_ vs. *τ*_2_ comparison, collapsed across intervals. Interval *p*-values are Greenhouse–Geisser corrected.

| Param. | Source | $F$ | df | $p$ | $\eta_p^2$ |
| --- | --- | --- | --- | --- | --- |
| TER | Group (control/forskolin) | 395.88 | 1, 10 | <0.001 | 0.975 |
|  | Interval | 287.48 | 3, 30 | 0.007 | 0.966 |
|  | Interaction | 279.85 | 3, 30 | <0.001 | 0.965 |
| TEC | Group (control/forskolin) | 11.88 | 1, 10 | 0.006 | 0.543 |
|  | Interval | 7.73 | 3, 30 | 0.032 | 0.436 |
|  | Interaction | 8.04 | 3, 30 | <0.001 | 0.446 |
| $\tau_1$ | Group (control/forskolin) | 1227.34 | 1, 10 | <0.001 | 0.992 |
|  | Interval | 843.93 | 3, 30 | 0.011 | 0.988 |
|  | Interaction | 1000.98 | 3, 30 | <0.001 | 0.990 |
| $\tau_2$ | Group (control/forskolin) | 578.22 | 1, 10 | <0.001 | 0.983 |
|  | Interval | 272.27 | 3, 30 | 0.004 | 0.965 |
|  | Interaction | 238.86 | 3, 30 | <0.001 | 0.960 |
| $\tau_1$ vs. $\tau_2$ | Group (control/forskolin) | 1021.86 | 1, 10 | <0.001 | 0.990 |
|  | Tau type | 1.13 | 1, 10 | 0.312 | 0.102 |
|  | Interaction | 12.06 | 1, 10 | 0.006 | 0.547 |

**A.6 Custom Chamber Design**

**Table A2:** Post-hoc pairwise comparisons for forskolin experiments (*n* = 6 per group). Independent *t*-tests were used for TER and TEC comparisons; paired *t*-tests were used for *τ*_1_ vs. *τ*_2_ comparisons. All corrections use the Holm–Bonferroni method.

| Param. | Comparison | Interval (min) | $t$ | $p_{\text{raw}}$ | $p_{\text{Holm}}$ |
| --- | --- | --- | --- | --- | --- |
| *TER: forskolin vs. control* |  |  |  |  |  |
| TER | forskolin vs. control | −10–0 | −1.24 | 0.244 | 0.244 |
| TER | forskolin vs. control | 0–10 | 13.93 | <0.001 | <0.001 |
| TER | forskolin vs. control | 10–20 | 20.76 | <0.001 | <0.001 |
| TER | forskolin vs. control | 20–30 | 21.69 | <0.001 | <0.001 |
| *TEC: forskolin vs. control* |  |  |  |  |  |
| TEC | forskolin vs. control | −10–0 | −0.16 | 0.877 | 0.877 |
| TEC | forskolin vs. control | 0–10 | −4.88 | <0.001 | 0.003 |
| TEC | forskolin vs. control | 10–20 | −2.34 | 0.041 | 0.099 |
| TEC | forskolin vs. control | 20–30 | −2.47 | 0.033 | 0.099 |
| *$\tau_1$ vs. $\tau_2$: control group* |  |  |  |  |  |
| $\tau_1$ vs. $\tau_2$ | control | −10–0 | 0.28 | 0.792 | 1.000 |
| $\tau_1$ vs. $\tau_2$ | control | 0–10 | 4.19 | 0.009 | 0.051 |
| $\tau_1$ vs. $\tau_2$ | control | 10–20 | 6.25 | 0.002 | 0.011 |
| $\tau_1$ vs. $\tau_2$ | control | 20–30 | 7.37 | <0.001 | 0.006 |
| *$\tau_1$ vs. $\tau_2$: FSK group* |  |  |  |  |  |
| $\tau_1$ vs. $\tau_2$ | forskolin | −10–0 | −0.12 | 0.909 | 1.000 |
| $\tau_1$ vs. $\tau_2$ | forskolin | 0–10 | −4.10 | 0.009 | 0.051 |
| $\tau_1$ vs. $\tau_2$ | forskolin | 10–20 | 0.28 | 0.788 | 1.000 |
| $\tau_1$ vs. $\tau_2$ | forskolin | 20–30 | 1.62 | 0.166 | 0.663 |

**Table A3:** ANOVA for simulated 20% decreases in *R_a_* and *R_b_* (*n* = 58 parameter sets). One-way repeated-measures ANOVA for TER and TEC; two-way repeated-measures ANOVA for parameter comparisons. Post-hoc paired *t*-tests with Holm–Bonferroni correction.

| Parameter | Source | $F$ | df | $p$ |
| --- | --- | --- | --- | --- |
| *Decreasing $R_b$* |  |  |  |  |
| TER | Interval | 1796.85 | 3, 171 | <0.001 |
| TEC | Interval | 0.38 | 3, 171 | 0.769 |
| $\tau_1$ vs. $\tau_2$ | Interval | 8650.54 | 3, 171 | <0.001 |
| | Term ($\tau$ type) | 8524.29 | 1, 57 | <0.001 |
|  | Interaction | 8807.27 | 3, 171 | <0.001 |
| $R_1$ vs. $R_2$ | Interval | 4873.01 | 3, 171 | <0.0001 |
| | Term ($R$ type) | 4746.81 | 1, 57 | <0.001 |
|  | Interaction | 4049.57 | 3, 171 | <0.001 |
| $C_1$ vs. $C_2$ | Interval | 854.81 | 3, 171 | <0.001 |
| | Term ($C$ type) | 892.33 | 1, 57 | <0.001 |
|  | Interaction | 885.92 | 3, 171 | <0.001 |
| *Decreasing $R_a$* |  |  |  |  |
| TER | Interval | 858.45 | 3, 171 | <0.001 |
| TEC | Interval | 1.87 | 3, 171 | 0.137 |
| $\tau_1$ vs. $\tau_2$ | Interval | 16488.80 | 3, 171 | <0.001 |
| | Term ($\tau$ type) | 417.15 | 1, 57 | <0.001 |
|  | Interaction | 428.20 | 3, 171 | <0.001 |
| $R_1$ vs. $R_2$ | Interval | 595.28 | 3, 171 | <0.001 |
| | Term ($R$ type) | 13274.04 | 1, 57 | <0.001 |
|  | Interaction | 12596.23 | 3, 171 | <0.001 |
| $C_1$ vs. $C_2$ | Interval | 4200.45 | 3, 171 | <0.001 |
| | Term ($C$ type) | 5378.56 | 1, 57 | <0.001 |
|  | Interaction | 5474.78 | 3, 171 | <0.001 |

**Table A4:**
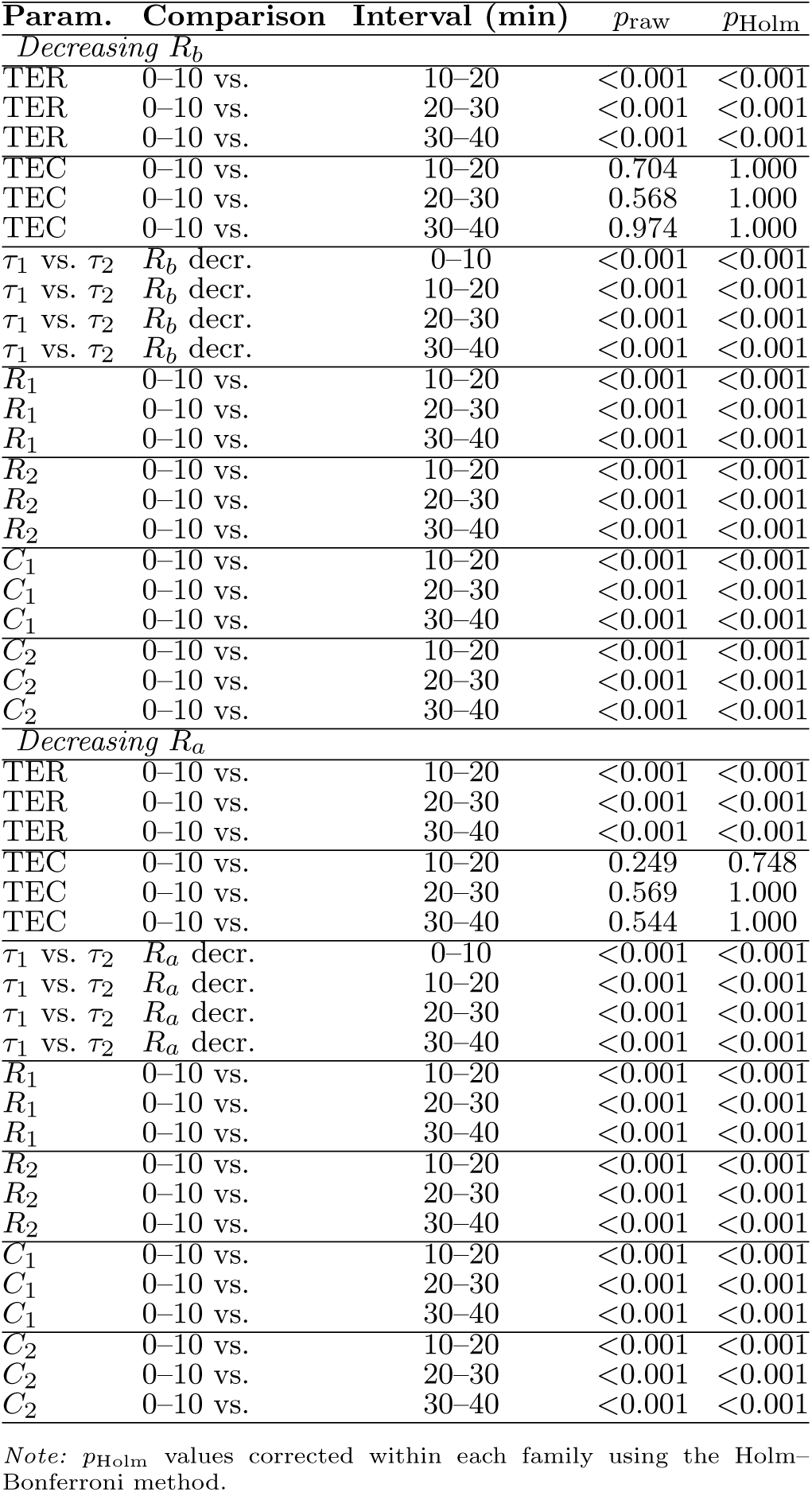
Post-hoc results for simulated 20% decreases in *R_a_* and *R_b_* (*n* = 58 parameter sets). One-way repeated-measures ANOVA for TER and TEC; two-way repeated-measures ANOVA for parameter comparisons. Post-hoc paired *t*-tests with Holm–Bonferroni correction.

| Param. | Comparison | Interval (min) | $p_{\text{raw}}$ | $p_{\text{Holm}}$ |
| --- | --- | --- | --- | --- |
| *Decreasing $R_b$* |  |  |  |  |
| TER | 0–10 vs. | 10–20 | <0.001 | <0.001 |
| TER | 0–10 vs. | 20–30 | <0.001 | <0.001 |
| TER | 0–10 vs. | 30–40 | <0.001 | <0.001 |
| TEC | 0–10 vs. | 10–20 | 0.704 | 1.000 |
| TEC | 0–10 vs. | 20–30 | 0.568 | 1.000 |
| TEC | 0–10 vs. | 30–40 | 0.974 | 1.000 |
| $\tau_1$ vs. $\tau_2$ | $R_b$ decr. | 0–10 | <0.001 | <0.001 |
| $\tau_1$ vs. $\tau_2$ | $R_b$ decr. | 10–20 | <0.001 | <0.001 |
| $\tau_1$ vs. $\tau_2$ | $R_b$ decr. | 20–30 | <0.001 | <0.001 |
| $\tau_1$ vs. $\tau_2$ | $R_b$ decr. | 30–40 | <0.001 | <0.001 |
| $R_1$ | 0–10 vs. | 10–20 | <0.001 | <0.001 |
| $R_1$ | 0–10 vs. | 20–30 | <0.001 | <0.001 |
| $R_1$ | 0–10 vs. | 30–40 | <0.001 | <0.001 |
| $R_2$ | 0–10 vs. | 10–20 | <0.001 | <0.001 |
| $R_2$ | 0–10 vs. | 20–30 | <0.001 | <0.001 |
| $R_2$ | 0–10 vs. | 30–40 | <0.001 | <0.001 |
| $C_1$ | 0–10 vs. | 10–20 | <0.001 | <0.001 |
| $C_1$ | 0–10 vs. | 20–30 | <0.001 | <0.001 |
| $C_1$ | 0–10 vs. | 30–40 | <0.001 | <0.001 |
| $C_2$ | 0–10 vs. | 10–20 | <0.001 | <0.001 |
| $C_2$ | 0–10 vs. | 20–30 | <0.001 | <0.001 |
| $C_2$ | 0–10 vs. | 30–40 | <0.001 | <0.001 |
| *Decreasing $R_a$* |  |  |  |  |
| TER | 0–10 vs. | 10–20 | <0.001 | <0.001 |
| TER | 0–10 vs. | 20–30 | <0.001 | <0.001 |
| TER | 0–10 vs. | 30–40 | <0.001 | <0.001 |
| TEC | 0–10 vs. | 10–20 | 0.249 | 0.748 |
| TEC | 0–10 vs. | 20–30 | 0.569 | 1.000 |
| TEC | 0–10 vs. | 30–40 | 0.544 | 1.000 |
| $\tau_1$ vs. $\tau_2$ | $R_a$ decr. | 0–10 | <0.001 | <0.001 |
| $\tau_1$ vs. $\tau_2$ | $R_a$ decr. | 10–20 | <0.001 | <0.001 |
| $\tau_1$ vs. $\tau_2$ | $R_a$ decr. | 20–30 | <0.001 | <0.001 |
| $\tau_1$ vs. $\tau_2$ | $R_a$ decr. | 30–40 | <0.001 | <0.001 |
| $R_1$ | 0–10 vs. | 10–20 | <0.001 | <0.001 |
| $R_1$ | 0–10 vs. | 20–30 | <0.001 | <0.001 |
| $R_1$ | 0–10 vs. | 30–40 | <0.001 | <0.001 |
| $R_2$ | 0–10 vs. | 10–20 | <0.001 | <0.001 |
| $R_2$ | 0–10 vs. | 20–30 | <0.001 | <0.001 |
| $R_2$ | 0–10 vs. | 30–40 | <0.001 | <0.001 |
| $C_1$ | 0–10 vs. | 10–20 | <0.001 | <0.001 |
| $C_1$ | 0–10 vs. | 20–30 | <0.001 | <0.001 |
| $C_1$ | 0–10 vs. | 30–40 | <0.001 | <0.001 |
| $C_2$ | 0–10 vs. | 10–20 | <0.001 | <0.001 |
| $C_2$ | 0–10 vs. | 20–30 | <0.001 | <0.001 |
| $C_2$ | 0–10 vs. | 30–40 | <0.001 | <0.001 |

*Note:* $p_{\text{Holm}}$ values corrected within each family using the Holm–Bonferroni method.

**Table A5:** ANOVA for simulated 20% increases in *R_a_* and *R_b_* (*n* = 58 parameter sets).

| Parameter | Source | $F$ | df | $p$ |
| --- | --- | --- | --- | --- |
| *Increasing $R_b$* |  |  |  |  |
| TER | Interval | 1906.62 | 3, 171 | <0.001 |
| TEC | Interval | 0.91 | 3, 171 | 0.438 |
| $\tau_1$ vs. $\tau_2$ | Interval | 5742.01 | 3, 171 | <0.001 |
| | Term ($\tau$ type) | 5942.00 | 1, 57 | <0.001 |
|  | Interaction | 5792.40 | 3, 171 | <0.001 |
| $R_1$ vs. $R_2$ | Interval | 3693.53 | 3, 171 | <0.001 |
| | Term ($R$ type) | 3745.03 | 1, 57 | <0.001 |
|  | Interaction | 3678.16 | 3, 171 | <0.001 |
| $C_1$ vs. $C_2$ | Interval | 921.90 | 3, 171 | <0.001 |
| | Term ($C$ type) | 954.11 | 1, 57 | <0.001 |
|  | Interaction | 957.88 | 3, 171 | <0.001 |
| *Increasing $R_a$* |  |  |  |  |
| TER | Interval | 878.21 | 3, 171 | <0.001 |
| TEC | Interval | 0.67 | 3, 171 | 0.573 |
| $\tau_1$ vs. $\tau_2$ | Interval | 10609.75 | 3, 171 | <0.001 |
| | Term ($\tau$ type) | 307.54 | 1, 57 | <0.001 |
|  | Interaction | 301.11 | 3, 171 | <0.001 |
| $R_1$ vs. $R_2$ | Interval | 657.71 | 3, 171 | <0.001 |
| | Term ($R$ type) | 21947.06 | 1, 57 | <0.001 |
|  | Interaction | 22658.15 | 3, 171 | <0.001 |
| $C_1$ vs. $C_2$ | Interval | 3589.41 | 3, 171 | <0.001 |
| | Term ($C$ type) | 4374.07 | 1, 57 | <0.001 |
|  | Interaction | 4292.44 | 3, 171 | <0.001 |

**Table A6:** Post-hoc results for simulated 20% increases in *R_a_* and *R_b_* (*n* = 58 parameter sets).

| Param. | Comparison | Interval | $p_{\text{raw}}$ | $p_{\text{Holm}}$ |
| --- | --- | --- | --- | --- |
| *Increasing $R_b$* |  |  |  |  |
| TER | 0–10 vs. | 10–20 | <0.001 | <0.001 |
| TER | 0–10 vs. | 20–30 | <0.001 | <0.001 |
| TER | 0–10 vs. | 30–40 | <0.001 | <0.001 |
| TEC | 0–10 vs. | 10–20 | 0.173 | 0.520 |
| TEC | 0–10 vs. | 20–30 | 0.659 | 1.000 |
| TEC | 0–10 vs. | 30–40 | 0.516 | 1.000 |
| $\tau_1$ vs. $\tau_2$ | $R_b$ incr. | 0–10 | <0.001 | <0.001 |
| $\tau_1$ vs. $\tau_2$ | $R_b$ incr. | 10–20 | <0.001 | <0.001 |
| $\tau_1$ vs. $\tau_2$ | $R_b$ incr. | 20–30 | <0.001 | <0.001 |
| $\tau_1$ vs. $\tau_2$ | $R_b$ incr. | 30–40 | <0.001 | <0.001 |
| $R_1$ | 0–10 vs. | 10–20 | <0.001 | <0.001 |
| $R_1$ | 0–10 vs. | 20–30 | <0.001 | <0.001 |
| $R_1$ | 0–10 vs. | 30–40 | <0.001 | <0.001 |
| $R_2$ | 0–10 vs. | 10–20 | <0.001 | <0.001 |
| $R_2$ | 0–10 vs. | 20–30 | <0.001 | <0.001 |
| $R_2$ | 0–10 vs. | 30–40 | <0.001 | <0.001 |
| $C_1$ | 0–10 vs. | 10–20 | <0.001 | <0.001 |
| $C_1$ | 0–10 vs. | 20–30 | <0.001 | <0.001 |
| $C_1$ | 0–10 vs. | 30–40 | <0.001 | <0.001 |
| $C_2$ | 0–10 vs. | 10–20 | <0.001 | <0.001 |
| $C_2$ | 0–10 vs. | 20–30 | <0.001 | <0.001 |
| $C_2$ | 0–10 vs. | 30–40 | <0.001 | <0.001 |
| *Increasing $R_a$* |  |  |  |  |
| TER | 0–10 vs. | 10–20 | <0.001 | <0.001 |
| TER | 0–10 vs. | 20–30 | <0.001 | <0.001 |
| TER | 0–10 vs. | 30–40 | <0.001 | <0.001 |
| TEC | 0–10 vs. | 10–20 | 0.433 | 1.000 |
| TEC | 0–10 vs. | 20–30 | 0.461 | 1.000 |
| TEC | 0–10 vs. | 30–40 | 0.782 | 1.000 |
| $\tau_1$ vs. $\tau_2$ | $R_a$ incr. | 0–10 | <0.001 | <0.001 |
| $\tau_1$ vs. $\tau_2$ | $R_a$ incr. | 10–20 | <0.001 | <0.001 |
| $\tau_1$ vs. $\tau_2$ | $R_a$ incr. | 20–30 | <0.001 | <0.001 |
| $\tau_1$ vs. $\tau_2$ | $R_a$ incr. | 30–40 | <0.001 | <0.001 |
| $R_1$ | 0–10 vs. | 10–20 | <0.001 | <0.001 |
| $R_1$ | 0–10 vs. | 20–30 | <0.001 | <0.001 |
| $R_1$ | 0–10 vs. | 30–40 | <0.001 | <0.001 |
| $R_2$ | 0–10 vs. | 10–20 | <0.001 | <0.001 |
| $R_2$ | 0–10 vs. | 20–30 | <0.001 | <0.001 |
| $R_2$ | 0–10 vs. | 30–40 | <0.001 | <0.001 |
| $C_1$ | 0–10 vs. | 10–20 | <0.001 | <0.001 |
| $C_1$ | 0–10 vs. | 20–30 | <0.001 | <0.001 |
| $C_1$ | 0–10 vs. | 30–40 | <0.001 | <0.001 |
| $C_2$ | 0–10 vs. | 10–20 | <0.001 | <0.001 |
| $C_2$ | 0–10 vs. | 20–30 | <0.001 | <0.001 |
| $C_2$ | 0–10 vs. | 30–40 | <0.001 | <0.001 |

*Note:* $p_{\text{Holm}}$ values corrected within each family using the Holm–Bonferroni method.

**Table A7:** Mixed ANOVA results for experimental perturbations. Forskolin: *n* = 6/group. GlyH-101: *n* = 6/group. Nystatin: *n* = 7/group. ETI/GlyH- 101: *n* = 6/group.

| Parameter | Source | $F$ | df | $p$ | $\eta_p^2$ |
| --- | --- | --- | --- | --- | --- |
| *GlyH-101* ($3 \times 4$ mixed ANOVA, $n = 6$) | | | | | |
| TER | Group | 21.16 | 2, 15 | <0.001 | 0.738 |
|  | Interval | 63.68 | 3, 45 | <0.001 | 0.809 |
|  | Interaction | 18.39 | 6, 45 | <0.001 | 0.710 |
| TEC | Group | 101.26 | 2, 15 | <0.001 | 0.931 |
|  | Interval | 129.47 | 3, 45 | <0.001 | 0.896 |
|  | Interaction | 31.86 | 6, 45 | <0.001 | 0.809 |
| *GlyH-101* — $\tau$ ($\tau \times \text{group} \times \text{Interval}$ factorial, $n = 6$) | | | | | |
| $\tau_1$ vs. $\tau_2$ | $\tau \times \text{group}$ | 6.45 | 2, 90 | 0.002 | — |
| | $\tau \times \text{Interval}$ | 0.004 | 2, 90 | 0.996 | — |
| | $\tau \times \text{group} \times \text{Interval}$ | 0.11 | 4, 90 | 0.980 | — |
| *Nystatin* ($2 \times 4$ mixed ANOVA, $n = 7$) | | | | | |
| TER | Group | 113.96 | 1, 12 | <0.001 | 0.905 |
|  | Interval | 153.42 | 3, 36 | <0.001 | 0.927 |
|  | Interaction | 49.32 | 3, 36 | <0.001 | 0.804 |
| TEC | Group | 0.25 | 1, 12 | 0.626 | 0.020 |
|  | Interval | 4.52 | 3, 36 | 0.038 | 0.274 |
|  | Interaction | 0.35 | 3, 36 | 0.790 | 0.028 |
| *Nystatin* — $\tau$ ($\tau \times \text{group} \times \text{Interval}$ factorial, $n = 7$) | | | | | |
| $\tau_1$ vs. $\tau_2$ | $\tau \times \text{group}$ | 12.31 | 1, 72 | <0.001 | — |
| | $\tau \times \text{Interval}$ | 0.75 | 2, 72 | 0.476 | — |
| | $\tau \times \text{group} \times \text{Interval}$ | 0.78 | 2, 72 | 0.463 | — |
| *ETI/GlyH-101* — TER ($2 \times 2 \times 4$ factorial, $n = 6$) | | | | | |
| Full factorial | Group × Interval | 63.41 | 3, 80 | <0.001 | — |
| | Group × ETI × Interval | 16.97 | 3, 80 | <0.001 | — |
| non-ETI | Group | 28.60 | 1, 10 | <0.001 | 0.741 |
|  | Interval | 61.73 | 3, 30 | 0.002 | 0.861 |
|  | Interaction | 34.31 | 3, 30 | <0.001 | 0.774 |
| ETI | Group | 116.98 | 1, 10 | <0.001 | 0.921 |
|  | Interval | 120.41 | 3, 30 | 0.005 | 0.923 |
|  | Interaction | 102.62 | 3, 30 | <0.001 | 0.911 |
| *ETI/GlyH-101* — TEC ($2 \times 2 \times 4$ factorial, $n = 6$) | | | | | |
| Full factorial | Group × Interval | 53.21 | 3, 80 | <0.001 | — |
| | Group × ETI × Interval | 34.71 | 3, 80 | <0.001 | — |
| non-ETI | Group | 8.29 | 1, 10 | 0.016 | 0.453 |
|  | Interval | 100.72 | 3, 30 | <0.001 | 0.910 |
|  | Interaction | 7.62 | 3, 30 | <0.001 | 0.433 |
| ETI | Group | 51.11 | 1, 10 | <0.001 | 0.836 |
|  | Interval | 39.74 | 3, 30 | 0.107 | 0.799 |
|  | Interaction | 134.66 | 3, 30 | <0.001 | 0.931 |
| *ETI/GlyH-101* — $\tau$ ($\tau \times \text{group} \times \text{ETI} \times \text{Interval}$ factorial, $n = 6$) | | | | | |
| Full factorial | $\tau \times \text{group}$ | 24.38 | 1, 120 | <0.001 | — |
| | $\tau \times \text{group} \times \text{ETI}$ | 14.80 | 1, 120 | <0.001 | — |
| | $\tau \times \text{group} \times \text{interval}$ | 15.81 | 2, 120 | <0.001 | — |
| | $\tau \times \text{group} \times \text{ETI} \times \text{interval}$ | 8.08 | 2, 120 | <0.001 | — |
| non-ETI | $\tau \times \text{group}$ | 1.06 | 1, 60 | 0.308 | — |
| | $\tau \times \text{group} \times \text{interval}$ | 0.05 | 2, 60 | 0.950 | — |
| ETI | $\tau \times \text{group}$ | 13.85 | 1, 60 | <0.001 | — |
| | $\tau \times \text{group} \times \text{Interval}$ | 8.98 | 2, 60 | <0.001 | — |

**Table A8:** Post-hoc pairwise comparisons for experimental perturbations. All with Holm– Bonferroni correction unless noted. CTL = control.

| Parameter | Comparison | Interval (min) | $n$ | $t$ | $p_{raw}$ | $p_{Holm}$ |
| --- | --- | --- | --- | --- | --- | --- |
| *GlyH-101 — TER, group main effect* |  |  |  |  |  |  |
| TER | CTL vs. 10 μM | — | 6 | −5.53 | <0.001 | <0.001 |
| TER | CTL vs. 5 μM | — | 6 | −9.27 | <0.001 | <0.001 |
| TER | 10 μM vs. 5 μM | — | 6 | 2.70 | 0.022 | 0.022 |
| *GlyH-101 — TER, interval × group* |  |  |  |  |  |  |
| TER | CTL vs. 10 μM | 0–10 | 6 | −5.49 | <0.001 | 0.002 |
| TER | CTL vs. 5 μM | 0–10 | 6 | −9.33 | <0.001 | <0.001 |
| TER | 10 μM vs. 5 μM | 0–10 | 6 | 2.90 | 0.016 | 0.095 |
| TER | CTL vs. 10 μM | 10–20 | 6 | −5.22 | <0.001 | 0.003 |
| TER | CTL vs. 5 μM | 10–20 | 6 | −8.87 | <0.001 | <0.001 |
| TER | 10 μM vs. 5 μM | 10–20 | 6 | 2.50 | 0.032 | 0.129 |
| TER | CTL vs. 10 μM | 20–30 | 6 | −5.66 | <0.001 | 0.002 |
| TER | CTL vs. 5 μM | 20–30 | 6 | −8.42 | <0.001 | <0.001 |
| TER | 10 μM vs. 5 μM | 20–30 | 6 | 2.61 | 0.026 | 0.129 |
| *GlyH-101 — TEC, group main effect* |  |  |  |  |  |  |
| TEC | CTL vs. 10 μM | — | 6 | 9.49 | <0.001 | <0.001 |
| TEC | CTL vs. 5 μM | — | 6 | 17.99 | <0.001 | <0.001 |
| TEC | 10 μM vs. 5 μM | — | 6 | 3.30 | 0.008 | 0.008 |
| *GlyH-101 — TEC, interval × group (selected)* |  |  |  |  |  |  |
| TEC | CTL vs. 10 μM | 0–10 | 6 | 9.11 | <0.001 | <0.001 |
| TEC | CTL vs. 5 μM | 0–10 | 6 | 15.96 | <0.001 | <0.001 |
| TEC | CTL vs. 10 μM | 10–20 | 6 | 8.12 | <0.001 | <0.001 |
| TEC | CTL vs. 5 μM | 10–20 | 6 | 19.41 | <0.001 | <0.001 |
| TEC | CTL vs. 10 μM | 20–30 | 6 | 7.71 | <0.001 | <0.001 |
| TEC | CTL vs. 5 μM | 20–30 | 6 | 19.46 | <0.001 | <0.001 |
| *GlyH-101 — $\tau_1$ vs. $\tau_2$ per group, collapsed across post-treatment intervals (paired $t$-test, $n = 6$)* |  |  |  |  |  |  |
| $\tau_1$ vs. $\tau_2$ | Control | 0–30 | 6 | 2.12 | 0.088 | 0.088 |
| $\tau_1$ vs. $\tau_2$ | 5 μM | 0–30 | 6 | −4.26 | 0.008 | 0.024 |
| $\tau_1$ vs. $\tau_2$ | 10 μM | 0–30 | 6 | −3.72 | 0.014 | 0.027 |
| *GlyH-101 — $T_{50}$ (Holm–Bonferroni across 4 tests)* |  |  |  |  |  |  |
| $T_{50}$: $\tau_1$ vs. $\tau_2$ | 10 μM | — | 6 | 9.30 | 0.0002 | 0.0010 |
| $T_{50}$: $\tau_2$ between doses | 5 vs. 10 μM | — | 6 | −6.97 | 0.0005 | 0.0017 |
| $T_{50}$: $\tau_1$ vs. $\tau_2$ | 5 μM | — | 6 | 7.57 | 0.0006 | 0.0013 |
| $T_{50}$: $\tau_1$ between doses | 5 vs. 10 μM | — | 6 | −0.72 | 0.489 | 0.489 |
| *Nystatin — TER (independent $t$-test, $n = 7$)* |  |  |  |  |  |  |
| TER | Nyst vs. CTL | −10–0 | 7 | −1.39 | 0.190 | 0.190 |
| TER | Nyst vs. CTL | 0–10 | 7 | 10.27 | <0.001 | <0.001 |
| TER | Nyst vs. CTL | 10–20 | 7 | 10.38 | <0.001 | <0.001 |
| TER | Nyst vs. CTL | 20–30 | 7 | 7.44 | <0.001 | <0.001 |
| *Nystatin — $T_{50}$: $\tau_1$ vs. $\tau_2$ (paired $t$-test, $n = 7$)* |  |  |  |  |  |  |
| $T_{50}$ | Nystatin | — | 7 | −6.45 | 0.0007 | — |
| *Nystatin — $\tau_1$ vs. $\tau_2$ per group, collapsed across post-treatment intervals (paired $t$-test, $n = 7$)* |  |  |  |  |  |  |
| $\tau_1$ vs. $\tau_2$ | Control | 0–30 | 7 | 2.58 | 0.042 | 0.042 |
| $\tau_1$ vs. $\tau_2$ | Nystatin | 0–30 | 7 | −19.68 | <0.001 | <0.001 |
| *ETI/GlyH-101 — TER, non-ETI (independent $t$-test, $n = 6$)* |  |  |  |  |  |  |
| TER | nonETI: GlyH vs. CTL | −10–0 | 6 | −0.17 | 0.865 | 0.865 |
| TER | nonETI: GlyH vs. CTL | 0–10 | 6 | −2.73 | 0.021 | 0.064 |
| TER | nonETI: GlyH vs. CTL | 10–20 | 6 | −5.21 | <0.001 | 0.002 |
| TER | nonETI: GlyH vs. CTL | 20–30 | 6 | −6.03 | <0.001 | <0.001 |
| *ETI/GlyH-101 — TER, ETI (independent $t$-test, $n = 6$)* |  |  |  |  |  |  |
| TER | ETI: GlyH vs. CTL | −10–0 | 6 | −1.59 | 0.144 | 0.287 |
| TER | ETI: GlyH vs. CTL | 0–10 | 6 | −10.47 | <0.001 | <0.001 |
| TER | ETI: GlyH vs. CTL | 10–20 | 6 | −11.11 | <0.001 | <0.001 |
| TER | ETI: GlyH vs. CTL | 20–30 | 6 | −10.38 | <0.001 | <0.001 |
| *ETI/GlyH-101 — TEC, non-ETI (independent $t$-test, $n = 6$)* |  |  |  |  |  |  |
| TEC | nonETI: GlyH vs. CTL | −10–0 | 6 | 0.63 | 0.542 | 0.542 |
| TEC | nonETI: GlyH vs. CTL | 0–10 | 6 | −3.55 | 0.005 | 0.022 |
| TEC | nonETI: GlyH vs. CTL | 10–20 | 6 | −3.65 | 0.004 | 0.022 |
| TEC | nonETI: GlyH vs. CTL | 20–30 | 6 | −2.18 | 0.054 | 0.162 |
| *ETI/GlyH-101 — TEC, ETI (independent $t$-test, $n = 6$)* |  |  |  |  |  |  |
| TEC | ETI: GlyH vs. CTL | −10–0 | 6 | −1.94 | 0.081 | 0.162 |
| TEC | ETI: GlyH vs. CTL | 0–10 | 6 | 9.19 | <0.001 | <0.001 |
| TEC | ETI: GlyH vs. CTL | 10–20 | 6 | 9.79 | <0.001 | <0.001 |
| TEC | ETI: GlyH vs. CTL | 20–30 | 6 | 8.08 | <0.001 | <0.001 |
| *ETI/GlyH-101 — $T_{50}$: $\tau_1$ vs. $\tau_2$ (paired $t$-test, $n = 6$)* |  |  |  |  |  |  |
| $T_{50}$ | ETI / glyh | — | 6 | 8.37 | <0.001 | — |
| *ETI/GlyH-101 — $\tau_1$ vs. $\tau_2$, ETI only (paired $t$-test, $n = 6$)* |  |  |  |  |  |  |
| $\tau_1$ vs. $\tau_2$ | ETI / CTL | 0–10 | 6 | 0.66 | 0.538 | 0.806 |
| $\tau_1$ vs. $\tau_2$ | ETI / CTL | 10–20 | 6 | 1.13 | 0.308 | 0.806 |
| $\tau_1$ vs. $\tau_2$ | ETI / CTL | 20–30 | 6 | 1.47 | 0.201 | 0.806 |
| $\tau_1$ vs. $\tau_2$ | ETI / GlyH | 0–10 | 6 | −18.31 | <0.001 | <0.001 |
| $\tau_1$ vs. $\tau_2$ | ETI / GlyH | 10–20 | 6 | 1.37 | 0.229 | 0.806 |
| $\tau_1$ vs. $\tau_2$ | ETI / GlyH | 20–30 | 6 | 1.98 | 0.104 | 0.522 |

Note: $p_{Holm}$ values corrected within each family using the Holm–Bonferroni method. Values $p < 0.05$ indicate statistical significance.

**Figure A1.**
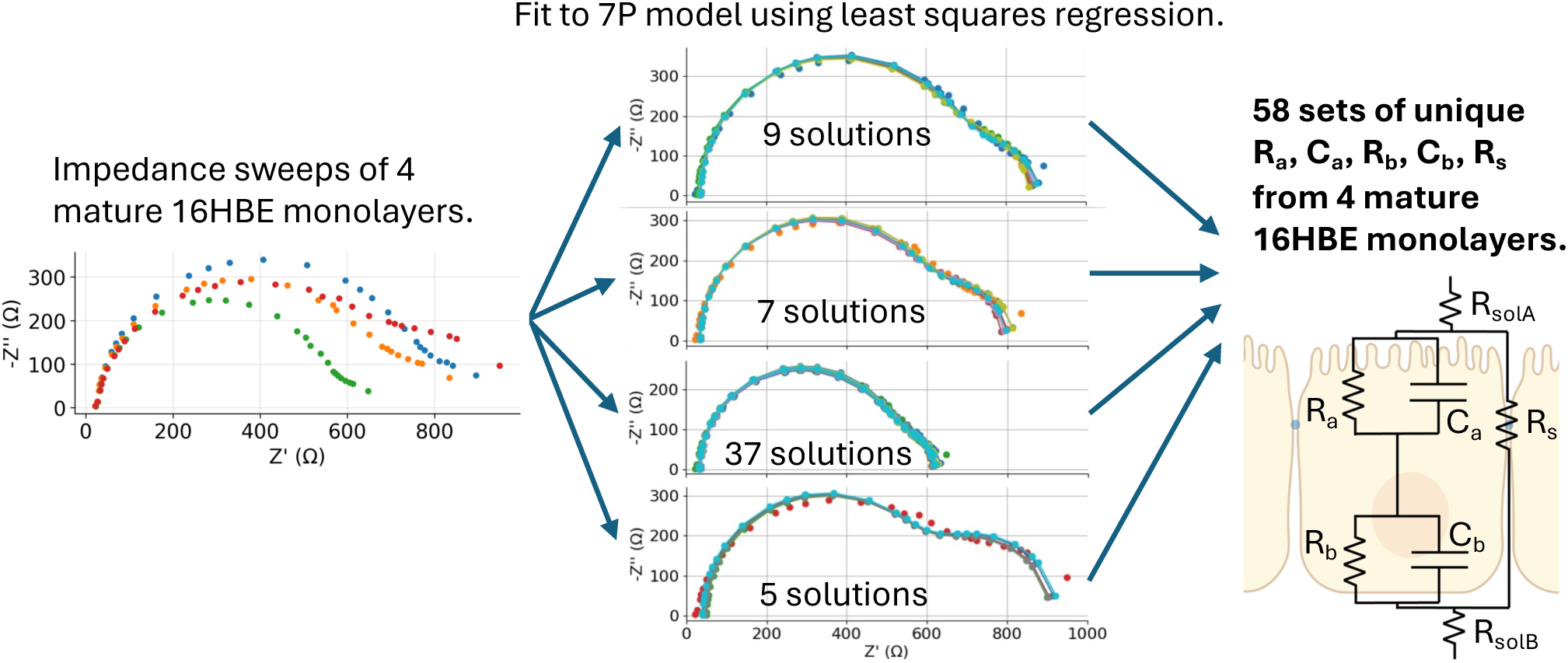
Method for determining physiologically relevant 7P initial values. Physiologically relevant 7P initial values were generated starting from impedance spectra of four mature 16HBE monolayers previously published [8]. Each impedance spectra is shown in the left Nyquist plot of negative imaginary impedance (-Z“) vs. real impedance (Z’). Multiple valid 7P solutions were obtained due to the underdetermined nature of the problem, resulting in 9, 7, 37, and 5 solutions for the four samples, each with unique 7P values (see Fig. A2), that served as the 58 initial values used in the simulated experiment.

**Figure A2.**
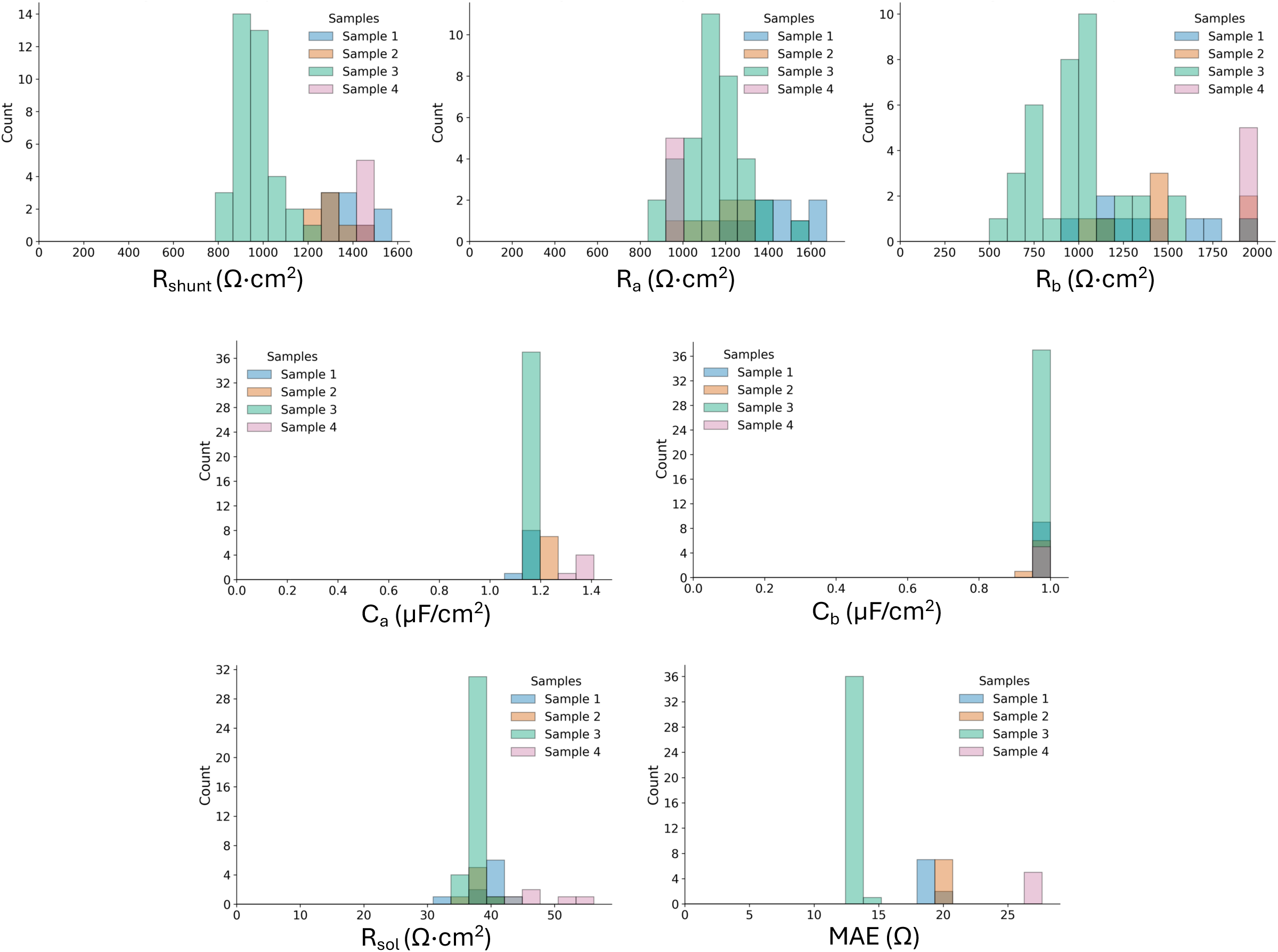
Least squares fitting of 16HBE impedance sweeps to the 7P model revealed several unique solutions within the solution space. 9, 7, 37, and 5 solutions for samples 1, 2, 3, and 4 (n=58 total). The local minima found have R_s_, R_a_, R_b_ varying several hundreds of Ω·cm^2^, but similar C_a_, C_b_, R_sol_ and mean absolute error (MAE, Ω) for each of the four biological samples.

**Figure A3.**
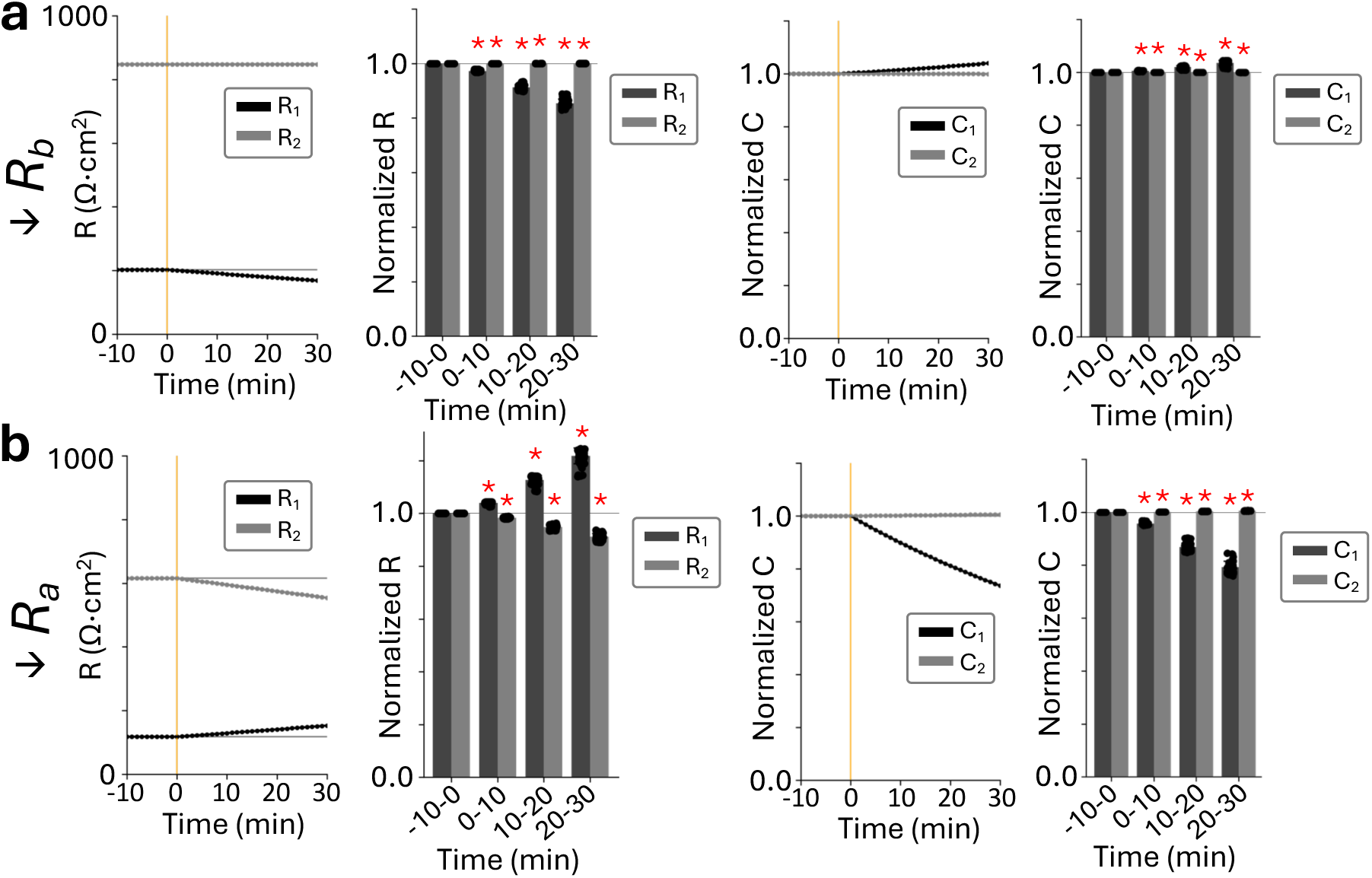
Effect of (a) R_b_ and (b) R_a_ decrease on individual RCRC parameters (n=58/group). (a) Repeated-measures ANOVA revealed significant main effects of time window and term, as well as significant interactions, for all parameters (R_1_, R_2_, C_1_, C_2_) under both (a) decreasing R_b_ and (b) decreasing R_a_ conditions (all p<0.001). Post-hoc paired t-tests with Holm correction confirmed that values at all subsequent time windows (0-10, 10–20, 20–30 min) differed significantly from the baseline-10-0 window for each parameter in both conditions (all corrected * p<0.001), but (a) decreasing R_b_ resulted in a 15.6% decrease in R_1_ with <1% change in R_2_ and a 3.5% increase in C_1_ with <1% change in C_2_ at the final interval. (b) Decreasing R_a_, resulted in an 8.7% decrease in R_2_. However, C_1_ also decreases and R_1_ increases 21.8%. These results suggest that changes in individual RCRC parameters do not map directly onto apical or basolateral resistance, even though the combined time constants **τ**_1_ and **τ**_2_ do.

**Figure A4.**
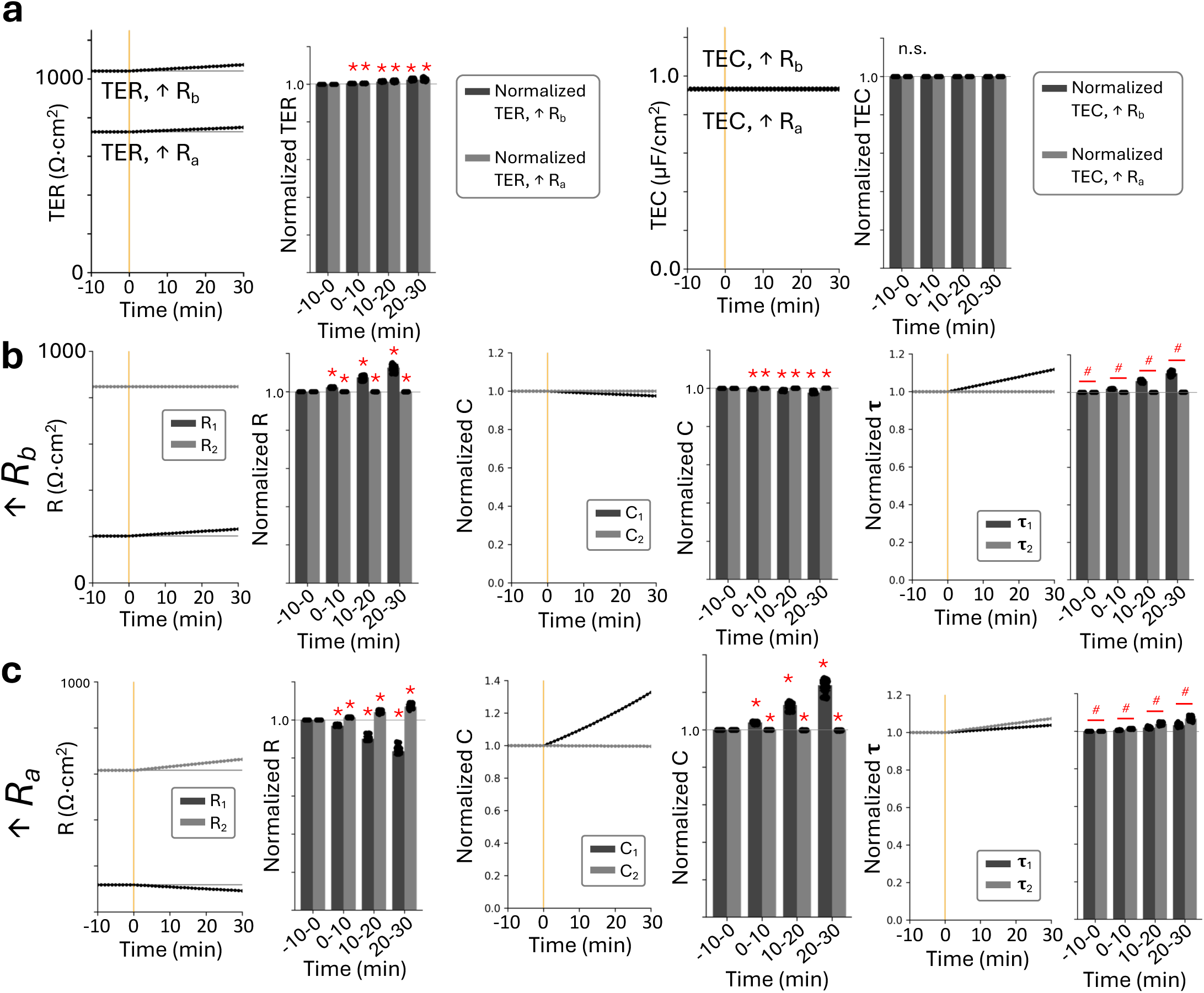
Effect of R_a_ and R_b_ increase on individual RCRC parameters (n=58/group). (a) Representative transepithelial resistance (TER) and transepithelial capacitance (TEC) simulated results and bar charts of the average value within each 10 min interval showed significant decrease over time (one-way repeated measures ANOVA, p<0.0001), with post-hoc paired t-tests (Holm-Bonferroni correction) confirming that all post-treatment intervals were significantly lower than the baseline interval (−10 to 0 min; *p<0.0001). TEC showed no significant change (n.s.) (b–c) Two-way repeated-measures ANOVA revealed significant main effects of time window and term, as well as significant interactions, for all RCRC parameters (R_1_, R_2_, C_1_, C_2_) under both (b) increasing R_b_ and (c) increasing R_a_ conditions (all p<0.001). Post-hoc paired t-tests (Holm-Bonferroni correction) confirmed that all parameters differed significantly from the baseline-10-0 min window at every subsequent interval (*p<0.001). (b) Increasing R_b_ produced a larger increase in R_1_ than R_2_, with minimal changes in C_1_ and C_2_. (c) Increasing R_a_ produced a larger increase in R_2_ but also caused a significant decrease in R_1_ and increase in C_1_. Despite these cross-parameter changes, **τ** values reliably separated the two conditions: increasing R_b_ produced a 10.1% increase in **τ**_1_ with <1% change in **τ**_2_, while increasing R_a_ produced a 6.7% increase in **τ**_2_ compared to only 3.2% in **τ**_1_ over 30 min. For both (b) and (c), a two-way repeated-measures ANOVA confirmed a significant τ × window interaction (p<0.001), with post-hoc paired t-tests (Holm-Bonferroni corrected) that showed **τ**_1_ significantly exceeded **τ**_2_ at all intervals (^#^p<0.001). These results demonstrated that individual RCRC parameters do not map directly onto apical or basolateral resistance, but the combined time constants **τ**_1_ and **τ**_2_ reliably distinguish between the two conditions. Complete ANOVA results and post-hoc tests are reported in Tables A3, A4.

**Figure A5.**
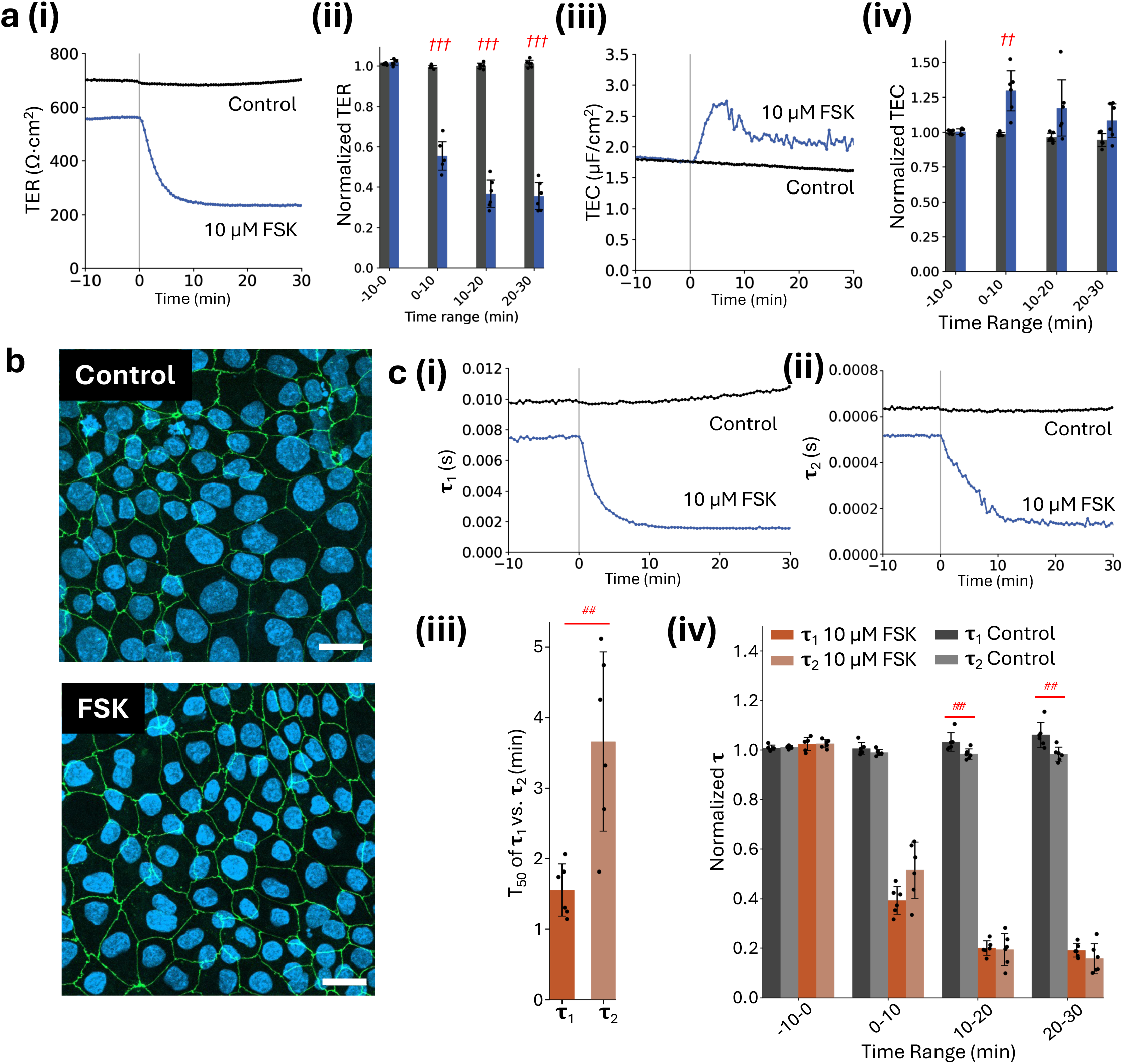
Effect of decreasing apical and basolateral membrane resistance with forskolin (FSK) on WT 16HBE (n=6/group). (a, i) Representative TER measurements showed that 10 µM forskolin produced a large, rapid TER decrease consistent with simultaneous increases in apical CFTR conductance and basolateral Na⁺–K⁺–2Cl⁻ cotransporter activity. TER fell by >44% relative to control. (a, ii) A 2 × 4 mixed ANOVA (group × time window) revealed significant main effects of group and time window, as well as a significant interaction, for both TER and TEC (all p<0.001). Post-hoc independent t-tests with Holm-Bonferroni correction showed no difference between control and forskolin groups at baseline (−10-0 min, p=0.244), but forskolin-treated samples were significantly lower than controls at all subsequent intervals (all corrected p<0.001). (a, iii-iv) Representative TEC measurements and bar charts show an immediate rise in response to forskolin, with TEC significantly higher than the control (28.0%, p<0.001), although this difference was transient as TEC decreases below 22% in the following intervals (n.s.) *^††^*p<0.01, *^†††^*p<0.001. (b) Following experiments representative maximum intensity Z-projection confocal images showed intact tight junctions for all groups; DAPI (blue), ZO-1 (green), scale bar 20 µm. (c, i-ii) Representative unnormalized **τ**_1_ and **τ**_2_ measurements. (c, iii) Half-maximal times (T_50_) indicated that **τ**_1_ responds faster than **τ**_2_ (mean difference 2.1 min, paired t-test, ^##^ p<0.01), consistent with rapid basolateral cotransporter activation. (c, iv) Forskolin treatment produced significant changes in both normalized **τ**_1_ and **τ**_2_ compared to controls. Mixed ANOVAs (group × window) revealed significant main effects and interactions for both **τ**_1_ and **τ**_2_ (both interaction p<0.001), with **τ**_1_ showing a stronger group-dependent response (η²_p_=0.992) than **τ**_2_ (η²_p_=0.983). A separate mixed ANOVA confirmed a significant group × **τ** interaction (p=0.006), indicating that forskolin differentially affected the two time constants. In the forskolin group, normalized **τ**_1_ and **τ**_2_ decreased rapidly following treatment, with **τ**_2_ significantly exceeding **τ**_1_ at the first post-treatment interval (**τ**_1_=0.39 vs. **τ**_2_=0.51, p=0.051), although this difference did not survive Holm–Bonferroni correction. At later intervals, **τ**_1_ and **τ**_2_ converged to similar amplitudes (all p>0.66), consistent with forskolin perturbing both membranes simultaneously. In the control group, **τ**_1_ and **τ**_2_ did not differ at baseline (-10-0 min, p=1.000) or in the 0-10 min interval (p=0.051), but **τ**_1_ became significantly greater than **τ**_2_ at the 10–20 and 20–30 min intervals (p=0.011 and p=0.006, respectively) ^#^ p<0.05, ^##^ p<0.01. Together, these results confirm that forskolin perturbs both membranes and that EEIS captures coordinated **τ**_1_ and **τ**_2_ responses when apical and basolateral permeabilities change simultaneously. This demonstrated apically administered compounds can elicit detectable basolateral changes, reinforcing the interpretation that the asymmetric **τ**_1_ and **τ**_2_ responses observed under selective perturbations reflect true membrane-specific dynamics rather than measurement artifacts.

**Figure A6.**
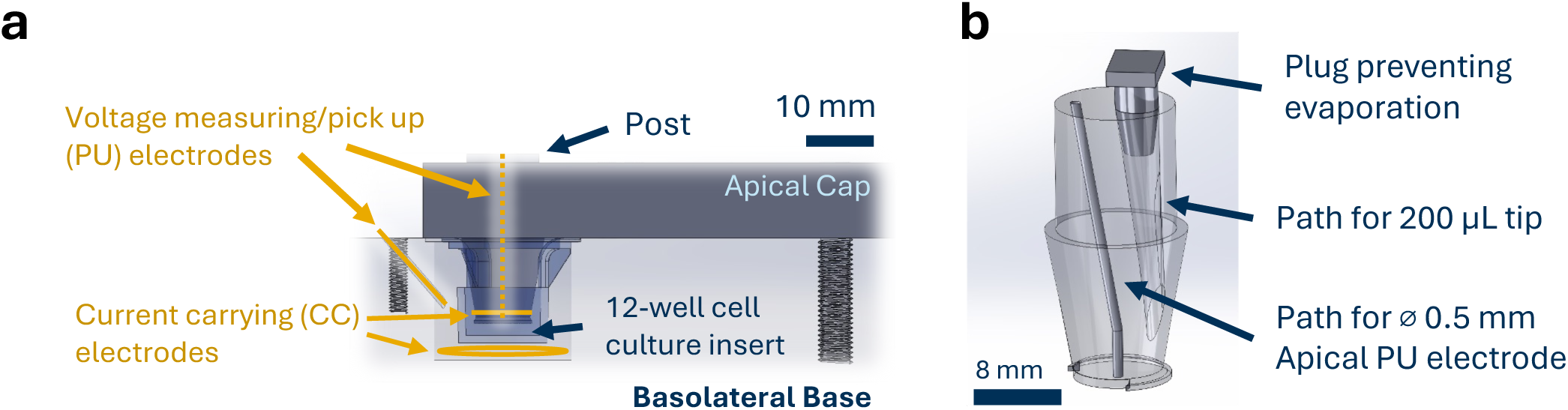
Custom Chamber Design. (A) Chamber milled from polycarbonate with 0.02’’ diameter 99.95% pure platinum wire (McMaster Carr, Elmhurst, IL) electrodes to enable repeatable voltage measuring and current carrying electrode placement. 15 mm diameter basolateral loop and 10 mm diameter apical loop as current sending electrodes, similar to previous designs [22] and straight wires with the tip submerged in the apical and basolateral baths will be used reference and sense voltage sensing electrodes. The wires were connected through alligator clips to the Autolab M204 galvanostat (Metrohm, Riverview, FL) to send current and measure voltage. (B) Central post enlarged, showing access point and path for addition of modulators during EEIS experiment, plug to prevent evaporation, and placement of apical voltage measuring electrode.

## References

[1] Anderson, J.M., Itallie, C.M.V.: Physiology and function of the tight junction. Cold Spring Harbor Perspectives in Biology 1, 2584–2585 (2009) 10.1101/CSHPERSPECT.A002584

[2] Cereijido, M., Robbins, E.S., Dolan, W.J., Rotunno, C.A., Sabatini, D.D.: Polarized monolayers formed by epithelial cells on a permeable and translucent support. The Journal of Cell Biology 77, 853 (1978) 10.1083/JCB.77.3.853

[3] Rizzolo, L.J., Peng, S., Luo, Y., Xiao, W.: Integration of tight junctions and claudins with the barrier functions of the retinal pigment epithelium. Progress in Retinal and Eye Research 30, 296–323 (2011) 10.1016/J.PRETEYERES.2011.06.002

[4] Singh, P., Arora, A., Strand, T.A., Leffler, D.A., Catassi, C., Green, P.H., Kelly, C.P., Ahuja, V., Makharia, G.K.: Global prevalence of celiac disease: Systematic review and meta-analysis. Clinical gastroenterology and hepatology 16, 823–8362 (2018) 10.1016/J.CGH.2017.06.037

[5] Eby, J.C., Ciesla, W.P., Hamman, W., Donato, G.M., Pickles, R.J., Hewlett, E.L., Lencer, W.I.: Selective translocation of the bordetella pertussis adenylate cyclase toxin across the basolateral membranes of polarized epithelial cells. Journal of Biological Chemistry 285, 10662–10670 (2010) 10.1074/jbc.M109.089219

[6] Ussing, H.H., Zerahn, K.: Active transport of sodium as the source of electric current in the short-circuited isolated frog skin. Acta Physiologica Scandinavica 23, 110–127 (1951) 10.1111/J.1748-1716.1951.TB00800.X

[7] Bialek, S., Miller, S.S.: K+ and cl-transport mechanisms in bovine pigment epithelium that could modulate subretinal space volume and composition. The Journal of Physiology 475, 401 (1994) 10.1113/JPHYSIOL.1994.SP020081

[8] Blaug, S., Quinn, R., Quong, J., Jalickee, S., Miller, S.S.: Retinal pigment epithelial function: a role for cftr? Documenta ophthalmologica. Advances in ophthalmology 106, 43–50 (2003) 10.1023/A:1022514031645

[9] Blaug, S., Hybiske, K., Cohn, J., Firestone, G.L., Machen, T.E., Miller, S.S.: Enac- and cftr-dependent ion and fluid transport in mammary epithelia. American Journal of Physiology - Cell Physiology 281 (2001) 10.1152/AJPCELL.2001.281.2.C633/ASSET/IMAGES/LARGE/H00810609015.JPEG

[10] Cotton, C.U., Stutts, M.J., Knowles, M.R., Gatzy, J.T., Boucher, R.C.: Abnormal apical cell membrane in cystic fibrosis respiratory epithelium. an in vitro electrophysiologic analysis. The Journal of Clinical Investigation 79, 80–85 (1987) 10.1172/JCI112812

[11] Hernandez, E.V., Hu, J.G., Frambach, D.A., Gallemore, R.P.: Potassium conductances in cultured bovine and human retinal pigment epithelium. Investigative Ophthalmology and Visual Science 36, 113–122 (1995)

[12] Hu, J.G., Gallemore, R.P., Bok, D., Frambach, D.A.: Chloride transport in cultured fetal human retinal pigment epithelium. Experimental Eye Research 62, 443–448 (1996) 10.1006/EXER.1996.0049

[13] Maminishkis, A., Chen, S., Jalickee, S., Banzon, T., Shi, G., Wang, F.E., Ehalt, T., Hammer, J.A., Miller, S.S.: Confluent monolayers of cultured human fetal retinal pigment epithelium exhibit morphology and physiology of native tissue. Investigative Ophthalmology & Visual Science 47, 3612–3624 (2006) 10.1167/IOVS.05-1622

[14] Quinn, R.H., Miller, S.S.: Ion transport mechanisms in native human retinal pigment epithelium. Investigative Ophthalmology and Visual Science 33, 3513– 3527 (1992)

[15] Rothenberg, P., Reuss, L., Glaser, L.: Serum and epidermal growth factor transiently depolarize quiescent bsc-1 epithelial cells. Proceedings of the National Academy of Sciences of the United States of America 79, 7783–7787 (1982) 10.1073/PNAS.79.24.7783;PAGEGROUP:STRING:PUBLICATION

[16] Tang, J., Abramcheck, F.J., Driessche, W.V., Helman, S.I.: Electro-physiology and noise analysis of k+-depolarized epithelia of frog skin. 10.1152/ajpcell.1985.249.5.C421 18 (1985) 10.1152/AJPCELL.1985.249.5.C421

[17] Welsh, M.J.: Anthracene-9-carboxylic acid inhibits an apical membrane, chloride conductance in canine tracheal epithelium. The Journal of Membrane Biology 78, 61–71 (1984) 10.1007/BF01872533/METRICS

[18] Miyagishima, K.J., Wan, Q., Corneo, B., Sharma, R., Lotfi, M.R., Boles, N.C., Hua, F., Maminishkis, A., Zhang, C., Blenkinsop, T., Khristov, V., Jha, B.S., Memon, O.S., D’Souza, S., Temple, S., Miller, S.S., Bharti, K.: In pursuit of authenticity: Induced pluripotent stem cell-derived retinal pigment epithelium for clinical applications. Stem Cells Translational Medicine 5, 1562 (2016) 10.5966/SCTM.2016-0037

[19] Kokkinaki, M., Sahibzada, N., Golestaneh, N.: Human induced pluripotent stem-derived retinal pigment epithelium (rpe) cells exhibit ion transport, membrane potential, polarized vascular endothelial growth factor secretion, and gene expression pattern similar to native rpe. Stem Cells 29, 825–835 (2011) 10.1002/STEM.635

[20] Lim, J.J., Fischbarg, J.: Electrical properties of rabbit corneal endothelium as determined from impedance measurements. Biophysical journal 36, 677–695 (1981) 10.1016/S0006-3495(81)84758-3

[21] Wills, N.K., Lewis, S.A., Eaton, D.C.: Active and passive properties of rabbit descending colon: A microelectrode and nystatin study. J. Membrane Biol 45, 81–108 (1979)

[22] Lewallen, C.F., Chien, A., Maminishkis, A., Hirday, R., Reichert, D., Sharma, R., Wan, Q., Bharti, K., Forest, C.R.: A biologically validated mathematical model for decoding epithelial apical, baso-lateral, and paracellular electrical properties. American journal of physiology: Cell physiology (2023) 10.1152/AJPCELL.00200.2023

[23] Chien, A.J., Lewallen, C.F., Khor, H., Cegla, A.V., Guo, R., Watson, A.L., Hatcher, C., McCarty, N.A., Bharti, K., Forest, C.R.: Method for extracellular electrochemical impedance spectroscopy on epithelial cell monolayers. Bio-protocol 15, 5341 10.21769/BIOPROTOC.5341

[24] Kreindler, J.L., Jackson, A.D., Kemp, P.A., Bridges, R.J., Danahay, H.: Inhibition of chloride secretion in human bronchial epithelial cells by cigarette smoke extract. American Journal of Physiology - Lung Cellular and Molecular Physiology 288, 894–902 (2005) 10.1152/AJPLUNG.00376.2004

[25] Clausen, C., Lewis, S.A., Diamond, J.M.: Impedance analysis of a tight epithelium using a distributed resistance model. Biophysical Journal 26, 291–317 (1979)

[26] Driessche, W.V., Vos, R.D., Jans, D., Simaels, J., Smet, P.D., Raskin, G.: Transepithelial capacitance decrease reveals closure of lateral interspace in a6 epithelia. Pflugers Archiv : European journal of physiology 437, 680–690 (1999) 10.1007/S004240050832

[27] Muller, S.M., Ebert, F., Bornhorst, J., Galla, H.-J., Francesconi, K.A., Schwerdtle, T.: Arsenic-containing hydrocarbons disrupt a model in vitro blood-cerebrospinal fluid barrier (2018) 10.1016/j.jtemb.2018.01.020

[28] Onnela, N., Savolainen, V., Juuti-Uusitalo, K., Vaajasaari, H., Skottman, H., Hyttinen, J.: Electric impedance of human embryonic stem cell-derived retinal pigment epithelium. Medical and Biological Engineering and Computing 50, 107– 116 (2012) 10.1007/S11517-011-0850-Z/TABLES/2

[29] Schifferdecker, E.: The ac impedance of necturus gallbladder epithelium. European Journal of Physiology 133, 125–133 (1978)

[30] Tosoni, K., Cassidy, D., Kerr, B., Land, S.C., Mehta, A.: Using drugs to probe the variability of trans-epithelial airway resistance. PLoS ONE 11, 0149550 (2016) 10.1371/JOURNAL.PONE.0149550

[31] Vazquez Cegla, A.J., Jones, K.T., Cui, G., Cottrill, K.A., Koval, M., McCarty, N.A.: Effects of hyperglycemia on airway epithelial barrier function in WT and CF 16HBE cells. Scientific reports 14(1), 25095 (2024) 10.1038/S41598-024-76526-3

[32] Gondzik, V., Awayda, M.S.: Methods for stable recording of short-circuit current in a na+-transporting epithelium. American Journal of Physiology: Cell Physiology 301, 162 (2011) 10.1152/ajpcell.00459.2010

[33] Sousa, F., Nascimento, C., Ferreira, D., Reis, S., Costa, P.: Reviving the interest in the versatile drug nystatin: A multitude of strategies to increase its potential as an effective and safe antifungal agent. Advanced Drug Delivery Reviews 199, 114969 (2023) 10.1016/J.ADDR.2023.114969

[34] Miller, S.S., Steinberg, R.H.: Passive ionic properties of frog retinal pigment epithelium. J. Membrane Biol 36, 337–372 (1977)

[35] Chien, A., Lewallen, C., Cui, G., Cegla, A.V., Lull, E., Khor, H., McCarty, N., Forest, C.: Rapid and accurate transepithelial resistance measurement of bronchiolar epithelium using impedance spectroscopy and rcrc model. Biophysical Journal

[36] Linz, G., Djeljadini, S., Steinbeck, L., Kose, G., Kiessling, F., Wessling, M.: Cell barrier characterization in transwell inserts by electrical impedance spectroscopy. Biosensors and Bioelectronics 165, 112345 (2020) 10.1016/J.BIOS.2020.112345

[37] Kottra, G., Fromter, E.: Rapid determination of intraepithelial resistance barriers by alternating current spectroscopy. i. experimental procedures. Pflugers Archiv: European journal of physiology 402, 409–420 (1984) 10.1007/BF00583942

[38] Kottra, G., Frömter, E.: Tight-junction tightness of necturus gall bladder epithelium is not regulated by camp or intracellular ca 2+ (ii. impedance measurements) 425, 535–545 (1993)

[39] Bello-Reuss, E.: Cell membranes and paracellular resistances in isolated renal proximal tubules from rabbit and ambystoma. The Journal of Physiology 370, 25 (1986) 10.1113/JPHYSIOL.1986.SP015920

[40] 16HBE14o-(WT CFTR) Cell Line Notes. https://www.cff.org/sites/default/files/2021-10/16HBE14o-Culturing-Protocol.pdf Accessed 2024-10-30

[41] Systematic optimization of prime editing for the efficient functional correction of cftr f508del in human airway epithelial cells. Nature Biomedical Engineering 2024 9:1 **9**, 7–21 (2024) 10.1038/s41551-024-01233-3

[42] Muanprasat, C., Sonawane, N.D., Salinas, D., Taddei, A., Galietta, L.J.V., Verkman, A.S.: Discovery of glycine hydrazide pore-occluding cftr inhibitors: mechanism, structure-activity analysis, and in vivo efficacy. Journal of general physiology 124, 125–137 (2004) 10.1085/jgp.200409059

[43] Cui, G., Cegla, A.V., Brown, J., Jones, K., Reed, R.C., Tirouvanziam, R., Koval, M., McCarty, N.A.: Hyperglycemia differentially affects neutrophil transmigration across cystic fibrosis and wildtype bronchial epithelia. Journal of Cystic Fibrosis 25, 127–134 (2026) 10.1016/j.jcf.2025.11.005

